# Elevated cortical tension triggers precocious ectopic polarization in mammalian oocytes, associated with lower developmental potential

**DOI:** 10.64898/2026.10.16.759562

**Authors:** A. Shihabi, E. Neiva, G. Letort, E. Nikalayevich, S. Madhavi, M. Yahiatene, X. Pollet-Villard, H. Turlier, MH. Verlhac, ME. Terret

**Author notes:** Correspondence (M-H.V), (M-E.T). These authors contributed equally (E. Neiva, G. Letort).

## Abstract

Oocytes from many organisms polarize early during development, unlike mammalian oocytes. The significance of maintaining this homogeneous state in mammals remains unclear. We show that elevated cortical tension, via increased actomyosin contractility or disrupted cortical architecture, induces precocious cortical polarization mid-meiosis I, with an ectopic actomyosin-rich domain opposite the future division site. This polarization generates directional cytoplasmic flows that displace organelles and the spindle, accelerating its migration and disrupting its orientation. A hydrodynamic model coupling the polarized cortex to a viscous cytoplasm accounts for these rearrangements. Ectopic polarization persists in meiosis II and is associated with mislocalization of fertilization factors, increased fertilization rates and polyspermy, and reduced early embryonic development. A similar morphological signature arises spontaneously in subsets of mouse and human oocytes. These results reveal precocious cortical polarization as a previously unrecognized, mechanically accessible state largely absent from mammalian oocytes, which may contribute to variability in oocyte quality.

## Introduction

Sexual reproduction in animals relies on the production of gametes through meiosis. In mammals, it ends with two successive divisions, meiosis I and II, that ultimately generate a single large haploid oocyte in females. A hallmark of oocyte divisions across the animal kingdom is their extreme asymmetry in the size of the resulting daughter cells. This precise geometry ensures that the organelles and maternal reserves accumulated during oocyte growth are retained predominantly in the oocyte’s cytoplasm, to be transmitted to the embryo after fertilization^1,2^. The geometry of cell division is dictated by the position of the microtubule spindle in the cell, and in oocytes, the asymmetric size of the division is therefore directly dependent on the cortical positioning of the spindle.

In many organisms, oocytes are polarized in Prophase prior to oocyte divisions, exhibiting an anisotropic organization of their cytoplasm and cortex, with the nucleus positioned at the cell cortex^3–12^. Upon entry into oocyte division, the spindle forms at the cortex where the nucleus was located, thereby promoting an asymmetric division. In contrast, mammalian oocytes lack polarization in Prophase, displaying neither cortical nor intracellular polarity^13–15^, and harbor a centrally positioned nucleus^16^. Consequently, after division entry, the spindle forms in the center of the oocyte and must migrate to the cortex to enable the asymmetry of the first division. As mammalian oocytes are not polarized in Prophase, the cortical site to which the spindle migrates is not predetermined; instead, the spindle axis forms randomly and the spindle migrates along its long axis toward the nearest cortex, a process that takes several hours^17^. Spindle migration is an actomyosin-dependent process, involving F-actin nucleators and force-generating elements, such as myosin-II, which mechanically pull the spindle toward the cortex^18–21^. This process also depends on the mechanical properties of the oocyte, particularly a decrease in cortical tension regulated by dynamic changes in actomyosin organization^20,22–24^. Once the spindle reaches the cortex at the end of meiosis I, it establishes oocyte polarity, characterized by an anisotropic organization of the cortex with a cortical actomyosin domain devoid of microvilli above the spindle (the cortical cap), while the remainder of the oocyte is covered with microvilli^25–27^. This cellular polarity is also present in meiosis II, as the spindle reforms beneath the cortex. The cortical cap is also depleted in CD9 and Juno, membrane proteins essential for sperm binding and fusion^28–30^. Accordingly, inhibiting oocyte polarity has deleterious effects on division and fertilization, as demonstrated by several studies in mice^13,31–34^. Thus, mechanical properties are critical for key aspects of oocyte development, including off-center spindle positioning at the cortex, which drives oocyte polarization at the end of meiosis I and enables the asymmetry of the first division^35^.

Cell mechanical properties regulate various biological processes, including proliferation, differentiation, migration and adhesion^36–39^. Therefore, their deregulation is involved in several human pathologies, such as cancer and aging^40–42^, and has recently been linked to embryo implantation failure^43^. Although oocytes do not proliferate, differentiate, migrate or adhere, the regulation of their mechanical properties is nevertheless essential for their developmental potential^20–22^. Interestingly, mechanical defects are relatively frequent in mammalian oocytes^44^, correlating with variability in cortical tension and in myosin-II cortical enrichment^20,22,24^. Notably, mouse and human oocytes with elevated cortical tension produce embryos with a reduced developmental potential^44^. Whether such mechanical defects remain confined to the cortex or propagate throughout the oocyte remains unknown.

To address the consequences of elevated cortical tension on oocyte development, we increased it using mechanistically distinct perturbations: constitutive RhoA activation at the cortex to enhance actomyosin contractility^24^, and alteration of cortical architecture resulting from the lack of the Arp2/3 complex in *arpc4-/-*oocytes^23,45^. Our results show that these perturbations converge on a common phenotype characterized by precocious cortical polarization mid-meiosis I, marked by the formation of a flattened, ectopic actomyosin enriched domain opposite the future division site. Strong directional cytoplasmic flows originate from this polarized domain, pushing both inert objects, such as oil droplets, and physiological structures, including the spindle and organelles, toward the opposite cortex. As a result, the spindle moves more rapidly and in a randomized orientation, consistent with quantitative analysis and biophysical modelling coupling cortical mechanics with cytoplasmic flows. These flows disrupt organelle distribution and reduce their inheritance after meiosis I, as measured for mitochondria and Golgi. During meiosis II, the ectopic polarization persists, leading to the spatially restricted accumulation of key fertilization components, including cortical granules, Juno, and CD9, within the polarized domain. This reorganization is associated with increased fertilization rates but also increased polyspermy. Consistently, early embryonic development of fertilized polarized oocytes is impaired. Finally, similar phenotypes are detected in natural populations of mouse and human oocytes using machine learning, supporting the potential physiological relevance of these observations. Together, these findings reveal that elevated cortical tension can drive a transition toward an aberrant polarized mechanical state that is normally absent in mammalian oocytes, resulting in large-scale cytoplasmic reorganization and altered fertilization and developmental outcomes.

## Results

### cRhoA oocytes polarize precociously mid-meiosis I, opposite to the future division site

Oocytes naturally vary in cortical tension and myosin-II cortical enrichment^20,22,24^. To elucidate the consequences of elevated cortical tension on oocyte development, we specifically enriched this subpopulation to be able to analyze a large number of cells. To achieve this, we developed a tool to increase cortical tension via the recruitment of active myosin-II at the cortex, using Ezrin as a cortical anchor as we previously described^21,22^. Specifically, we adapted a construct^46^ encoding a fusion protein, termed cRhoA (for cortical RhoA), comprising the RhoA(GEF) domain of LARG (a nucleotide exchange factor of the RhoA GTPase), Ezrin (which stably localizes at the oocyte cortex^47^), and mCherry for detection (Fig S1A). cRhoA is recruited at the cortex via Ezrin, promoting local RhoA-GTP production and activation of the RhoA/ROCK pathway^48^, which in turn phosphorylates Myosin Light Chain (MLC) and activates myosin-II^46^. We confirmed by live imaging and immunofluorescence that cRhoA localizes to the cortex in Prophase I (Fig S1B), and induces the recruitment of active myosin-II (p-MLC2 in Fig S1C, quantification of p-MLC2 cortical enrichment in Fig S1D), compared to control oocytes (either non-injected oocytes, or those expressing a fusion protein of Ezrin and mCherry). Micropipette aspiration and Atomic Force Microscopy from a previous study demonstrated that cRhoA oocytes have increased cortical tension in Prophase I^24^, consistent with a more circular shape (aspect ratio closer to 1) than controls (Fig S1E). Notably, expression of Ezrin alone does not affect cortical tension nor oocyte development^22^.

Having validated our tool to increase cortical tension via the recruitment of active myosin-II at the cortex, we assessed the development of cRhoA oocytes compared to controls using live microscopy (Fig 1). cRhoA oocytes divided properly (Fig 1A, Movie 1) in terms of timing (Fig 1B), efficiency (Fig 1C) and geometry (Fig 1D), extruding polar bodies comparable in surface area to those of controls (Fig 1E). However, upon entry into meiosis I, they formed multiple cortical zones enriched in cRhoA that progressively merged into a single, large, well defined polarized domain by mid-meiosis I (Fig 1F, pink asterisk). This polarized region was also enriched in actin and active myosin-II, as shown by immunofluorescence (Fig S1F). In contrast, Ezrin did not polarize in control oocytes (Fig 1F). This precocious polarization occurred in virtually all oocytes expressing cRhoA (Fig 1G). To gain insight into the dynamics of cRhoA polarization, we employed optimal transport techniques (drawing on methodologies similar to those described in^49^) to track the distribution of cRhoA pixel intensities in the cortex in consecutive frames, allowing to determine the displacement field of the cortex over time (see Supplementary Methods). This enabled qualitative and quantitative analysis of dynamic spatial changes of cRhoA fluorescence in time at the oocyte cortex, as shown in the averaged cRhoA velocity kymograph (Fig 1H). Small zones enriched in cRhoA were present as of nuclear envelope breakdown (BD), and merged to form a large single one (centered around 180° Fig 1H) from mid-meiosis I onward, always opposite the future division site (located at 0°/360° Fig 1H). We then plotted the average polarization speed from the averaged cRhoA velocity kymographs (Fig 1I). The merging of the cRhoA small zones was progressive with an increase in speed from BD until mid-meiosis I when a single stable region was formed. This analysis of cortical flows is robust, as the same pattern was found in movies analyzed at different time resolutions (Fig S2A-D). The same analysis was done for control oocytes expressing Ezrin (Fig S2E-F), and as expected, there was no displacement of Ezrin fluorescence in time and space at the cortex.

**Figure 1:**
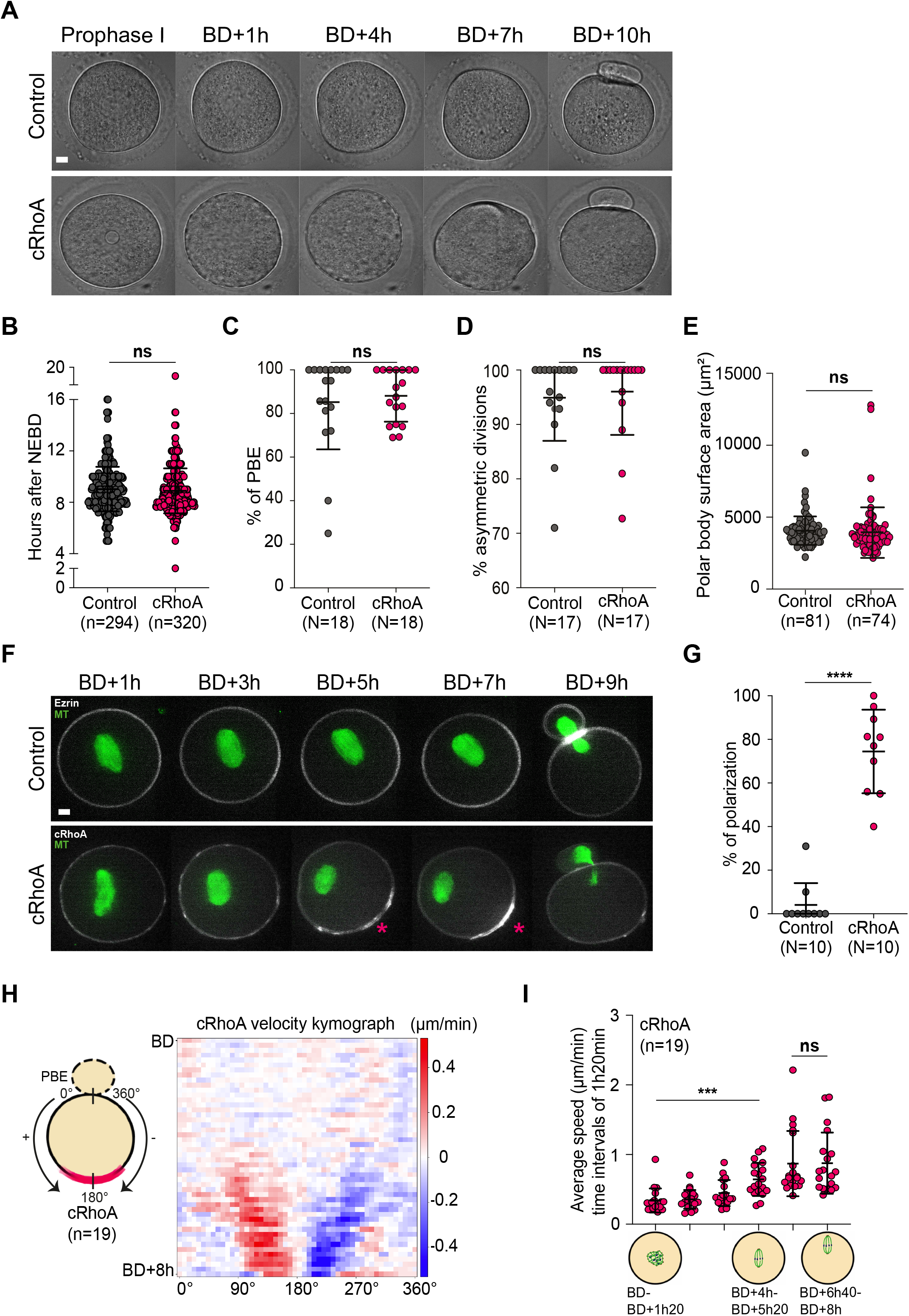
cRhoA oocytes divide normally but polarize precociously mid-meiosis I. (A) Time-lapse transmitted light images of control and cRhoA oocytes between Prophase I and BD+10h. (B) Timing of PBE (in hours after NEBD) in control and cRhoA oocytes. Data from 21 independent experiments. Means ± SD are shown, each dot is 1 oocyte. Statistical significance of differences was assessed by a Mann-Whitney test, P = 0.1044. (C) Percentage of control and cRhoA oocytes that did PBE. Data from 18 independent experiments. Means ± SD are shown, each dot is the percentage from one experiment. 273 control and 267 cRhoA oocytes were analyzed. Statistical significance of differences was assessed by a Mann-Whitney test, P = 0.8391. (D) Percentage of control and cRhoA oocytes that divided asymmetrically. Data from 17 independent experiments. Means ± SD are shown, each dot is the percentage from one experiment. 257 control and 246 cRhoA oocytes were analyzed. Statistical significance of differences was assessed by a Mann-Whitney test, P = 0.3875. (E) Polar body surface area (µm^2^) of control and cRhoA oocytes. Data from 14 independent experiments. Means ± SD are shown, each dot is 1 oocyte. Statistical significance of differences was assessed by a Mann-Whitney test, P = 0.1156. (F) Time-lapse fluorescent images of control and cRhoA oocytes between BD+1h and BD+9h. Microtubules (MT, green) were visualized with SiR-Tubulin. Control oocytes express Ezrin (white), cRhoA oocytes express cRhoA (white), which progressively accumulates (pink asterisk) opposite to where polar body extrusion will take place. Images are 12 µm Z-projections. (G) Percentage of control and cRhoA oocytes that polarized during meiosis I. Data from 10 independent experiments. Means ± SD are shown, each dot is the percentage from one experiment. 154 control and 156 cRhoA oocytes were analyzed. Statistical significance of differences was assessed by a Mann-Whitney test, P < 0.0001. (H) Left: scheme showing the orientation of the oocyte cortex for analysis (0° and 360°: where the polar body is extruded; 180°: cRhoA polarization). Right: Averaged cRhoA velocity kymograph at the cortex between BD and PBE-10 min, computed from 19 films with images acquired every 10 min, collected from 3 independent experiments. The X-axis represents the unfolded oocyte cortex as shown on the scheme (left), and the Y-axis represents the time from BD until BD+8h and the velocity of cRhoA fluorescence (µm/min). The motion of pixel intensities is positive between 0° and 180° and represented in red, negative between 360° and 180° and represented in blue, and the colors are more vivid at higher speeds. (I) Average speed of polarization of cRhoA at the cortex between BD and BD+8h using the data from the averaged cRhoA velocity kymograph in H. Each dot is 1 oocyte. Data from 19 films, collected from 3 independent experiments. Statistical significance of differences was assessed by a Wilcoxon matched-pairs signed rank test, from left to right P = 0.0003, P = 0.7381. For all graphs, cRhoA oocytes are in pink, controls in grey, n is the number of oocytes, N is the number of experiments. For all images, scale bar: 10 µm. BD or NEBD: Nuclear envelope breakdown; PBE: Polar Body Extrusion.

Overall, our data show that oocytes with increased cortical tension in Prophase I polarize precociously mid-meiosis I, with an ectopic region enriched in actomyosin opposite to the future division site. Interestingly, elevated cortical tension drives cortical instability and actomyosin polarization in other models, such as breast cancer cells^50^ and zebrafish embryonic cells^51^, switching their migration mode to adapt to their environment. However, oocytes are non-motile cells, so it is unlikely that polarization would induce a net movement of the cell. We therefore asked whether it could induce movements within the cell.

### Actomyosin polarization in cRhoA oocytes generates aberrant cytoplasmic flows that perturb spindle migration and the cytoplasmic landscape

We quantified cytoplasmic movements using spatiotemporal image correlation spectroscopy (STICS) analysis (see Methods), focusing from mid-meiosis I onward, when the precocious ectopic polarization is established in cRhoA oocytes, until polar body extrusion (Fig 2A-B, Movie 2). This analysis determines the direction and velocity of cytoplasmic particles by extracting velocity vectors from temporal and spatial correlations^16,52^ (see Methods). From mid-meiosis I onward, cRhoA oocytes exhibited stronger (Fig 2A-B) and more directional (Fig 2A) cytoplasmic flows along the cortex, directed toward the polarized domain and subsequently toward the future site of the first division. Before mid-meiosis I, cytoplasmic flows were comparable in cRhoA and control oocytes (Fig S3A-B, Movie 2). Notably, the cortex of cRhoA oocytes flattened at the site of polarization, altering the aspect ratio of these oocytes, which became more oval compared to controls (Fig S3D-E). This suggests that actomyosin accumulation could induce a local contraction of the cortex^53,54^ which in turn would push the cytoplasm and generate directional cytoplasmic flows originating from the polarized region away from it.

**Figure 2:**
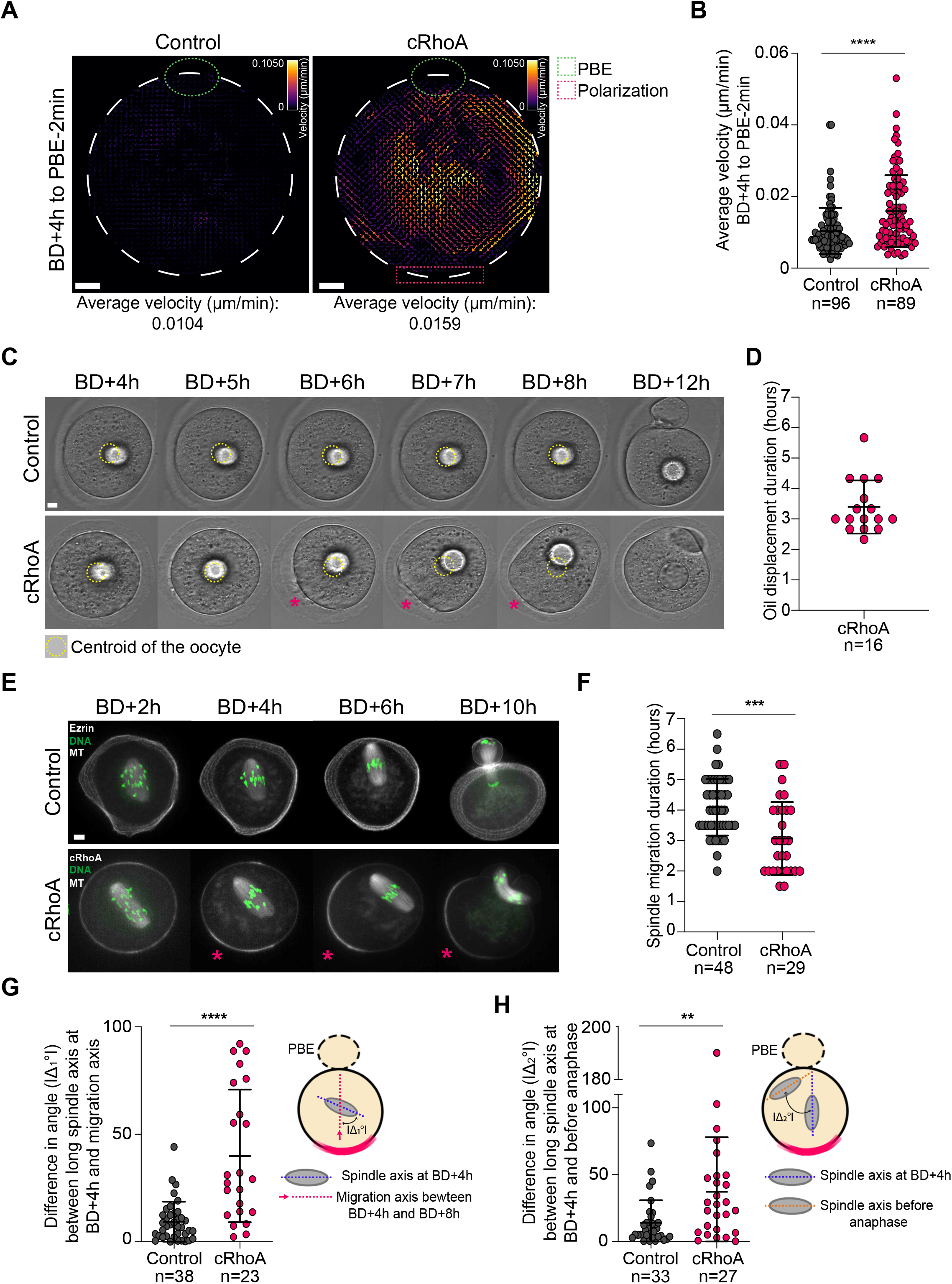
Strong flows emerge from the polarized region toward the future division site in cRhoA oocytes, pushing objects away from it and impacting spindle migration. (A) Representative vector maps of cytoplasmic flows from transmitted light videos obtained using STICS analysis for control and cRhoA oocytes, measured between BD+4h and PBE-2 min. (B) Average velocity, calculated from all average velocity vectors between BD+4h and PBE-2 min, for control and cRhoA oocytes. Data from 14 independent experiments. Means ± SD are shown, each dot is 1 oocyte. Statistical significance of differences was assessed by a Mann-Whitney test, P < 0.0001. (C) Time-lapse transmitted light images of control and cRhoA oocytes injected with an oil droplet from BD+4h until BD+12h. The yellow dotted circle shows the centroid of the oocyte at each time point during meiosis I. The pink asterisk indicates where cRhoA oocytes polarize. (D) Duration of oil droplet displacement (hours) in meiosis I for cRhoA oocytes. Data from 5 independent experiments. Mean ± SD is shown. (E) Time-lapse fluorescent images of control and cRhoA oocytes between BD+2h and BD+10h. Microtubules (MT, white) were visualized with SPY-Tubulin and DNA (green) was visualized with SPY-DNA. Control oocytes express Ezrin (white), cRhoA oocytes express cRhoA (white), which progressively accumulates (pink asterisk) opposite to where polar body extrusion will take place. Images are 12 to 15 µm Z-projections. (F) Duration of spindle migration (hours) in meiosis I for control and cRhoA oocytes. Data from 6 independent experiments. Mean ± SD is shown. Statistical significance of differences was assessed by a Mann-Whitney test, P = 0.0002. (G) Left: absolute difference in angle (|Δ_1_°|) between the long axis of the spindle at BD+4h and the axis of spindle migration. Right: scheme illustrating this metric. Data from 6 independent experiments. Mean ± SD is shown. Statistical significance of differences was assessed by a Mann-Whitney test, P < 0.0001. (H) Left: absolute difference in angle (|Δ_2_°|) between the long axis of the spindle at BD+4h and just before anaphase. Right: scheme illustrating this metric. Data from 6 independent experiments. Mean ± SD is shown. Statistical significance of differences was assessed by a Mann-Whitney test, P = 0.0049. For all graphs, cRhoA oocytes are in pink, controls in grey, n is the number of oocytes. For all images, scale bar: 10 µm. BD: Nuclear envelope breakdown; PBE: Polar Body Extrusion.

We next tested whether these directional cytoplasmic flows originating from the polarized region were strong enough to displace objects within the cytoplasm. To this end, we injected oil droplets in the cytoplasm in Prophase I and followed their movement until polar body extrusion (Fig 2C, Movie 3). As previously shown, oil droplets remained centered in control oocytes^55^. In contrast, in cRhoA oocytes, oil droplets were pushed away from the flattened polarized region toward the future site of the first division, over a period of 3 hours and 20 minutes (Fig 2C-D). These results indicated that the cytoplasmic flows are sufficient to displace large passive objects in the cytoplasm, suggesting that they could affect other physiological structures comparable in size. To test this, we monitored spindle displacement in cRhoA oocytes in live throughout meiosis I (Fig 2E, Movie 4). We first computed the time it took for spindles to reach the cortex in control and cRhoA oocytes, starting from mid-meiosis I when the spindles were bipolar located at the oocyte center, and when cRhoA oocytes were polarized. In cRhoA oocytes, the spindle reached the cortex faster than in controls, in about 3 hours (Fig 2F), comparable to the duration of displacement of the oil droplets. This suggests that these movements are powered by the same mechanism, different from the one that normally off-centers the spindle in control oocytes. We then analyzed the characteristics of spindle displacement in control and cRhoA oocytes. We measured the absolute difference in angle (|Δ_1_°|) between the long axis of the spindle at the beginning of spindle migration, and the migration axis (Fig 2G). In controls, the spindle migrates to the closest cortex along its long axis^17^, thus the difference in angle between the long axis of the spindle and the migration axis is close to zero (Fig 2G). In cRhoA oocytes, the difference in angle between the long axis of the spindle and the migration axis was very variable, from zero to almost 100°. It suggested that some cRhoA oocytes resembled control oocytes, with the spindle migrating along its long axis, whereas in many cRhoA oocytes, the spindle migrated at an angle, sometimes even with its long axis parallel to the cortex (Fig 2E, G). We then assessed the persistence of spindle movement by measuring the absolute difference in angle (|Δ_2_°|) between the long axis of the spindle at the beginning and at the end of spindle migration (Fig 2H). In controls, the spindle migrates along its long axis in a persistent manner^17^, thus the difference in angle between the long axis of the spindle at the beginning and end of spindle migration is close to zero (Fig 2H). In cRhoA oocytes, the difference in angle between the long axis of the spindle at the beginning and end of spindle migration was very variable, from zero to almost 200°, showing that the spindle of some cRhoA oocytes rotated during migration (Fig 2H).

To better understand the physical mechanisms underlying the observed phenomena in cRhoA oocytes, we developed a biophysical model of cytoplasmic streaming based on coupled cortex-cytoplasm viscous flows (Fig 3A). The numerical model builds on a recent framework for modelling coupled surface-bulk viscous flows in animal cells^56^ with governing equations derived from a simple model of cellular symmetry breaking^57^ and discretized using the Finite Element (FE) method^58^. We adapted the model^56^ to represent the spindle and its movement (see Supplementary Methods). In this model, the cortex is described as an active fluid where myosin-II-dependent surface tension gradients generate Marangoni-like stresses that drive cortical flows, while the underlying cytoplasm is modeled as a passive viscous fluid advected through no-slip coupling at the interface, resulting in a fountain flow pattern (Fig 3A). The model was solved numerically under an axisymmetric approximation (see Supplementary Methods). To simplify computations, the spindle was approximated as a rigid, frictionless permeable body that did not perturb the flow, allowing its translation and rotation to be computed from the simulated velocity field. The model was calibrated with quantitative experimental measurements of spatiotemporal changes in oocyte geometry, obtained from transmitted light images covering the entire duration of the physical process. Analysis of these data revealed a traveling cortical contraction band characterized by equatorial contraction and polar expansion (Fig 3B). The model quantitatively reproduced this pattern by fitting a space-and time-dependent law for cortical myosin-II concentration, governing active tension dynamics (Fig 3B). The model was then validated by comparing its predicted flow fields with time-averaged flow fields from STICS analyses, which showed good agreement in directionality (Fig 3C). Importantly, the model predicted that cytoplasmic streaming exerts hydrodynamic torques on the spindle depending on its position relative to the oocyte center, leading to robust spindle rotation across a broad range of initial conditions (Fig 3D-E, Movie 5). This behavior aligns with experimental measurements of spindle dynamics, particularly regarding the rotation sense of the spindle (Fig 3F), supporting a mechanism in which myosin-II-driven cortical flows organize cytoplasmic streaming and cell-scale deformations while simultaneously moving the spindle through flow-induced torques.

**Figure 3:**
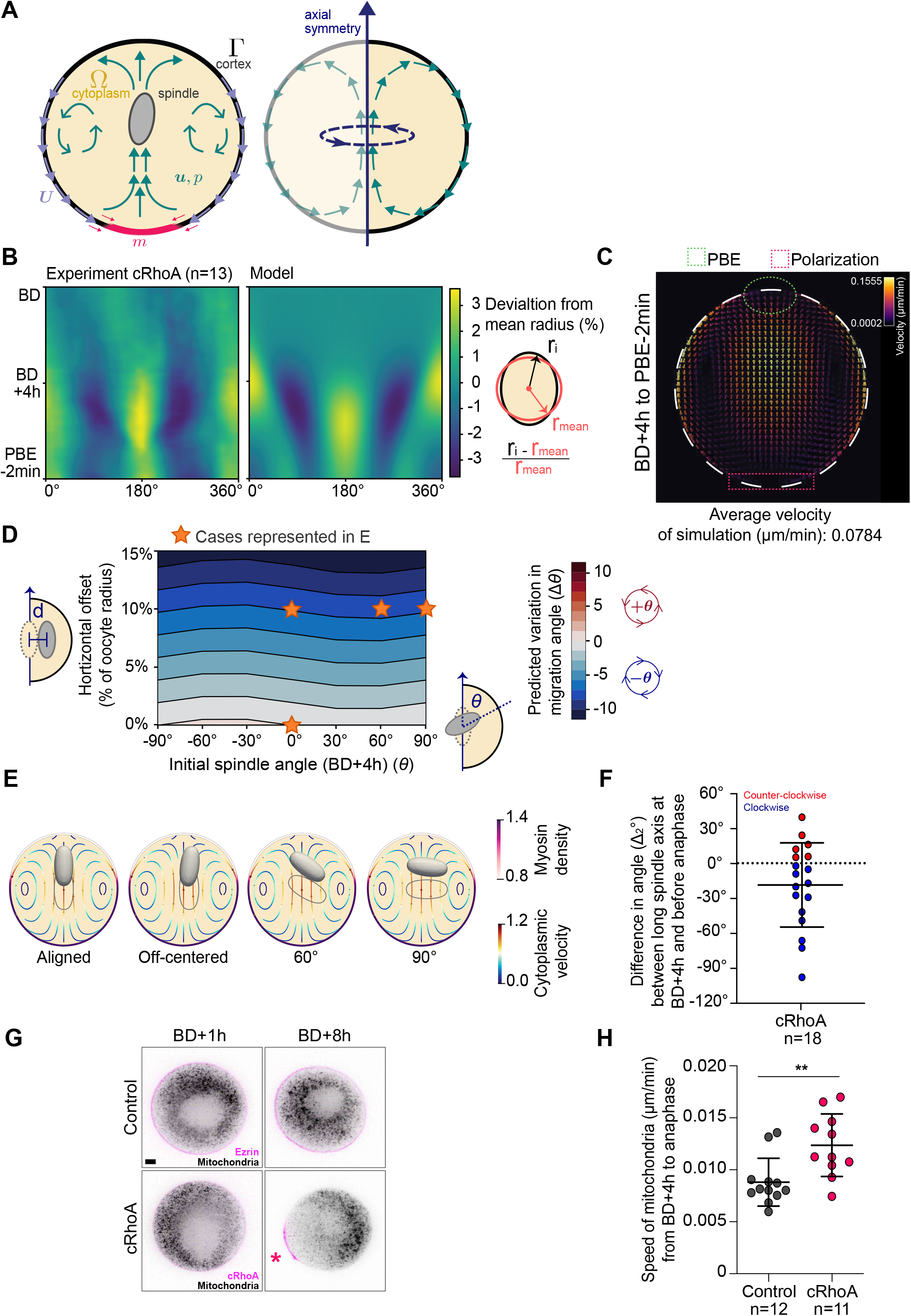
A hydrodynamic model of cytoplasmic streaming explains spindle migration defects in cRhoA oocytes, which also exhibit altered mitochondria distribution. (A) Schematic of the model. The cortex is represented as an active viscous surface in which myosin-II-dependent tension gradients generate cortical flows that advect the underlying cytoplasm, modelled as a passive viscous fluid. Coupling between the two phases produces a fountain flow pattern. (B) Kymographs of oocyte radius deviation from the mean radius in experiments (left panel) and simulations (right panel), revealing a travelling cortical contraction band from the equator toward the cRhoA-enriched region. The model reproduces the spatiotemporal pattern and amplitude of contraction and expansion. (C) Simulated cytoplasmic velocity field. (D) Model predictions of spindle rotation across the parameter space defined by initial spindle orientation and horizontal offset, showing robust rotation driven by cytoplasmic flows. (E) Representative projections from simulations illustrating spindle dynamics for different initial configurations, highlighting the dependence of rotational behavior on initial conditions. The spindle’s initial position is indicated only by its outline, while its final position is indicated by its outline filled with gray. (F) Experimental measurements of spindle rotation in cRhoA oocytes, showing a preferential rotation sense consistent with model predictions. Each dot is 1 oocyte. Data from 3 independent experiments. (G) Fluorescent images of control and cRhoA oocytes at BD+1h and BD+8h. Mitochondria (black) were visualized with Mitotracker. Control oocytes express Ezrin (pink), cRhoA oocytes express cRhoA (pink), which progressively accumulates (pink asterisk) opposite to where polar body extrusion will take place. Images are 8 to 12 µm Z-projections. Scale bar: 10 µm. BD: Nuclear envelope breakdown. (H) Speed of mitochondria (µm/min) from BD+4h to anaphase in control and cRhoA oocytes obtained using PIV analysis. Data was gathered from 7 independent experiments. Mean ± SD is shown, each dot is 1 oocyte. Statistical significance of differences was assessed by a Mann-Whitney test, P = 0.0045. BD: Nuclear envelope breakdown.

Our results show that cytoplasmic flows in cRhoA oocytes can displace large objects in the cytoplasm, such as the spindle. We assessed whether smaller objects, such as mitochondria, could also be displaced by these flows. We monitored mitochondria in cRhoA oocytes by live imaging throughout meiosis I (Fig 3G, Movie 6). Mitochondria localized similarly in cRhoA and control oocytes in early meiosis I, being distributed throughout the cytoplasm (Fig 3G). However, in cRhoA oocytes, mitochondria were progressively displaced away from the polarization zone, accumulating toward the future division site, whereas they aggregated around the spindle in controls^15^ (Fig 3G, Movie 6). We quantified mitochondria movement using Particle Image Velocimetry (PIV, see Methods), and showed that mitochondria moved faster in cRhoA oocytes (Fig 3H), in the same range as cytoplasmic flow velocities (Fig 2B). This suggests that mitochondria are pushed by the cytoplasmic flows originating from the polarized region in cRhoA oocytes.

To exclude the possibility that the phenotypes observed in cRhoA oocytes result from cRhoA expression itself rather than from increased cortical tension, we sought to examine an independent model of elevated cortical tension. We previously showed that the physiological decrease in cortical tension during meiosis I relies on Arp2/3-dependent actin nucleation at the cortex, which promotes the exclusion of cortical myosin-II^20,22–24^. We therefore analyzed oocytes lacking Arp2/3-dependent actin nucleation due to the conditional inactivation of the Arp2/3 subunit ArpC4 (*arpc4-/-*)^23^. Micropipette aspiration measurements confirmed that *arpc4-/-*oocytes display elevated cortical tension compared with controls (Fig S4A). Strikingly and similarly to cRhoA oocytes, *arpc4-/-*oocytes underwent precocious polarization mid-meiosis I (Fig S4B, pink asterisk), leading to the emergence of directional cytoplasmic flows comparable to those present in cRhoA oocytes (Fig S4C, Movie 2). Consistent with our observations in cRhoA oocytes, spindle migration was also affected in *arpc4-/-*oocytes, with the spindle frequently changing orientation during migration to the cortex (Fig S4D-E). Because *arpc4-/-*oocytes display additional early alterations affecting organelle organization and dynamics^23^, we did not further compare organelle-related phenotypes between the two models.

Altogether, our data show that elevated cortical tension, regardless of whether it arises from increased actomyosin contractility (cRhoA oocytes) or altered cortical architecture (*arpc4-/-*oocytes), is sufficient to trigger precocious oocyte polarization mid-meiosis I, forming an ectopic actomyosin-enriched zone from which cytoplasmic flows emerge, propelling objects such as the spindle and organelles toward the future division site. We went on to analyze cRhoA oocytes after the first meiotic division, in meiosis II.

### cRhoA oocytes are ectopically polarized in meiosis II, with mislocalization of fertilization factors associated with abnormal fertilization and embryonic development

First, we investigated whether cRhoA oocytes were polarized in meiosis II using live microscopy (Fig S5A, Movie 7). Strikingly, cRhoA oocytes displayed two polarized regions at the end of meiosis I, which persisted into meiosis II (Fig S5A, pink and green asterisks, similar to *arpc4-/-*oocytes in meiosis I Fig S4B). The first one was the polarized region enriched in cRhoA (Fig S5A, pink asterisk). The second one, located on the opposite side, was the cortical cap (Fig S5A, green asterisk). The cortical cap was also present in control oocytes (Fig S5A, green asterisk). This is a region enriched in actomyosin and devoid of microvilli, CD9 and Juno, membrane proteins involved in sperm adhesion and fusion (Fig 4A-B), and it is induced by the proximity of the spindle bearing the chromosomes to the cortex^25,27–29,59^. In cRhoA oocytes, CD9 and Juno accumulated in the cRhoA-rich polarized region, an area that was much smaller than in control oocytes (Fig 4A-B). Nonetheless, sperm were able to attach to the oocyte on this restricted region (Fig S5B). In addition, cortical granules, whose exocytosis following sperm fusion prevents polyspermy^60^, were also accumulated in the cRhoA-rich polarized region (Fig S5C).

**Figure 4:**
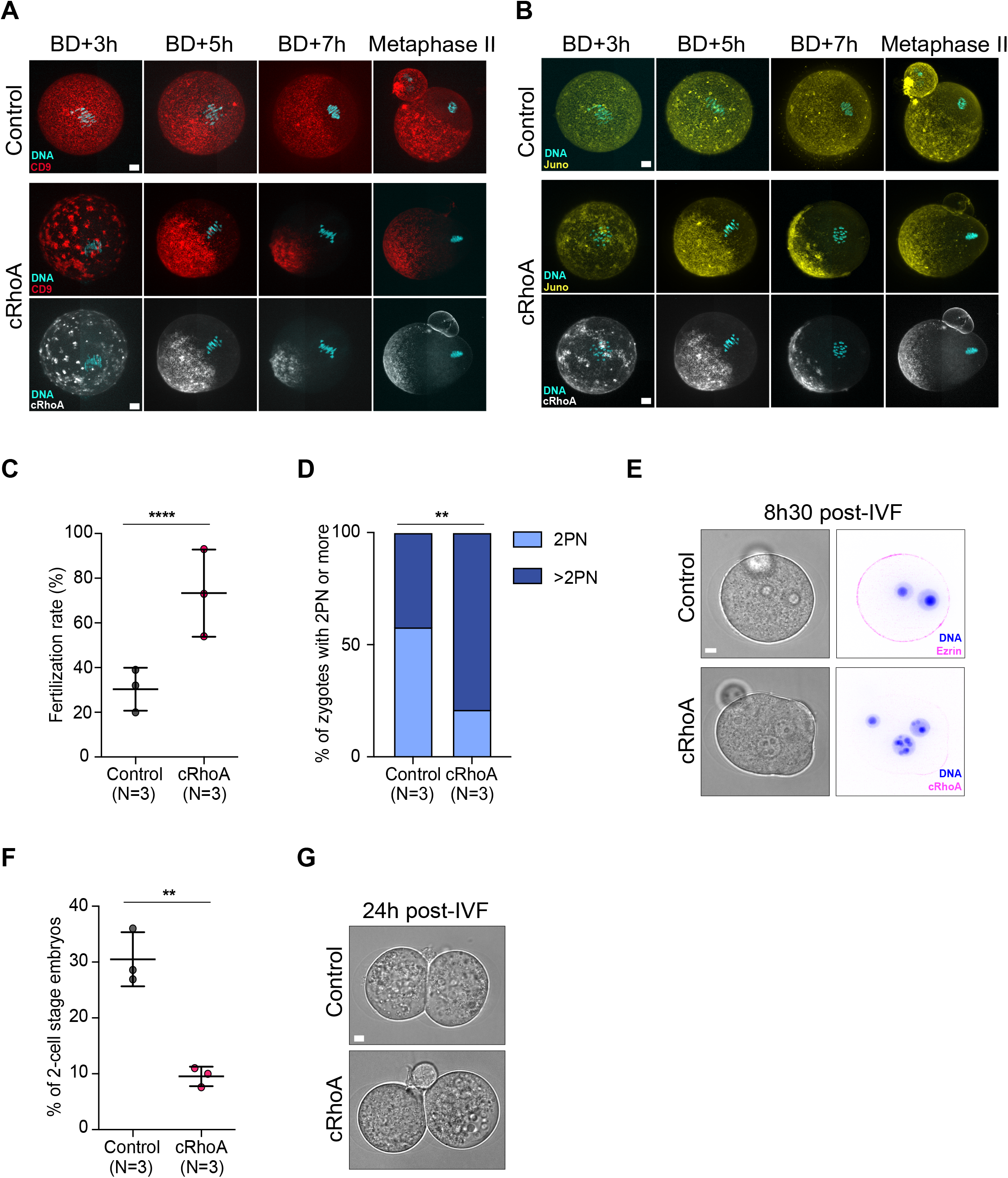
Fertilization factors are mislocalized in cRhoA oocytes during meiosis I and II, associated with abnormal fertilization and embryonic development. (A) Fluorescent images of control and cRhoA oocytes between BD+3h and Metaphase II. DNA (blue) was visualized with SPY-DNA and CD9 (red) with an FITC α-mouse CD9 monoclonal antibody. cRhoA oocytes express cRhoA (white, bottom). Images are Z-projections of the entire oocyte. (B) Time-lapse fluorescent images of control and cRhoA oocytes between BD+3h and Metaphase II (same oocytes as in A for some timepoints). DNA (blue) was visualized with SPY-DNA and Juno (yellow) with an AlexaFluor 647 α-mouse FR4 monoclonal antibody. cRhoA oocytes express cRhoA (white, bottom). Images are Z-projections of the entire oocyte. (C) Percentage of control and cRhoA oocytes fertilized after *in vitro* fertilization (IVF, displaying at least two pronuclei, including normally fertilized and polyspermic zygotes). Data from 3 independent experiments. Means ± SD are shown, each dot is the percentage from one experiment. 78 control and 68 cRhoA oocytes were analyzed. Statistical significance of differences was assessed by an exact Cochran-Mantel-Haenszel test stratified by experiment, P < 0.0001. (D) Percentage of fertilized control and cRhoA oocytes (zygotes) with 2 pronuclei (PN, light blue) or more than 2PN (dark blue). Data from 3 independent experiments. 23 control and 51 cRhoA zygotes were analyzed. Statistical significance of differences was assessed by an exact Cochran-Mantel-Haenszel test stratified by experiment, comparing the number of zygotes with 2PN versus more than 2PN, P = 0.005. (E) Transmitted light and fluorescent images of control and cRhoA zygotes 8h30 post-IVF. Control zygotes express Ezrin (pink), cRhoA zygotes express cRhoA (pink). DNA is in blue (H2B-GFP). One Z plane is shown for Ezrin and cRhoA, and a 28 µm Z-projection for the DNA. (F) Percentage of control and cRhoA oocytes that reached the 2-cell stage after IVF. Data from 3 independent experiments. Means ± SD are shown, each dot is the percentage from one experiment. 77 control and 51 cRhoA embryos were analyzed. Statistical significance of differences was assessed by an exact Cochran-Mantel-Haenszel test stratified by experiment, P = 0.005. (G) Transmitted light images of control and cRhoA 2-cell stage embryos 24h post-IVF. For all graphs, cRhoA oocytes are in pink, controls in grey, N is the number of independent experiments. For all images, scale bar: 10 µm. BD: Nuclear envelope breakdown. PN: pronuclei. IVF: *in vitro* fertilization.

In control oocytes in meiosis II, actin flows emerge from the cortical cap inducing cytoplasmic streaming and a pushing force on the spindle, maintaining it at the cortex^52^. We therefore asked whether these flows were altered in cRhoA oocytes. We quantified cytoplasmic movements in meiosis II using STICS analysis (Fig S3A, C, Movie 2). cRhoA oocytes displayed directional (Fig S3A) cytoplasmic flows from the cRhoA-rich polarized region away from it toward the cortical cap. These flows were stronger (Fig S3A, C) but oriented in the same direction as those in the controls^52^ (Fig S3A). As in meiosis I, they were able to push mitochondria away from the cRhoA-rich polarized region (Fig S5D). Interestingly, cRhoA oocytes displayed lower mitochondria and Golgi content in meiosis II compared to controls (Fig S5D-G).

Given that cRhoA oocytes exhibit abnormal localization of CD9, Juno and cortical granules, as well as a non-canonical cytoplasmic landscape, we assessed their fertilization competence and early developmental potential. Both control and cRhoA oocytes were matured *in vitro* and subjected to *in vitro* fertilization (IVF) after removal of their zona pellucida. Fertilization and polyspermy were evaluated by counting pronuclei in zygotes. The fertilization rate was defined as the percentage of zygotes displaying at least two pronuclei (including normally fertilized and polyspermic zygotes). Because zona-free oocytes generally display high fertilization rates and increased susceptibility to polyspermy, *in vitro* fertilization experiments were performed under conditions designed to maintain low fertilization rates and moderate polyspermy in control oocytes, thereby enabling detection of differences between experimental groups (Fig 4C-D, see Methods). Notably, a higher proportion of cRhoA oocytes were fertilized compared with controls (Fig 4C, 69.7% versus 30.3%). However, among fertilized oocytes, the proportion of zygotes with exactly two pronuclei was markedly lower in cRhoA oocytes than in controls (Fig 4D-E, 21.2% versus 56.5% in controls), consistent with increased polyspermy. In a separate set of experiments, only 9.5% of cRhoA oocytes that underwent IVF progressed to the 2-cell stage, compared with 30.5% of control oocytes (Fig 4F-G), indicating reduced developmental potential.

Together, our data reveal that cRhoA oocytes are aberrantly polarized during meiosis II, associated with the mislocalization of fertilization factors and a non-canonical cytoplasmic organization. This altered architecture correlates with increased sperm incorporation but defective polyspermy control and reduced early embryonic development progression.

### A subset of mouse and human oocytes naturally display the phenotype of cRhoA oocytes

Since cRhoA oocytes have been modified to artificially increase their cortical tension in Prophase I, we next asked whether their phenotype exists in natural populations of mouse and human oocytes, given that some of them exhibit spontaneously elevated cortical tension and myosin-II cortical enrichment^20,22,24,44^. We used our machine learning pipeline Oocytor^23,61,62^ to extract morphological and dynamic features from transmitted light movies of control and cRhoA oocytes progressing through meiosis I, using polar body extrusion (PBE) as a temporal reference (Fig 5A). Only oocytes in which PBE occurred within the imaging plane were included. A Random Forest classifier trained on these features revealed that discrimination between control and cRhoA oocytes varied over time, with a peak approximately 40 minutes before PBE (Fig 5B), which indicated that the morphological differences between the two populations were pronounced at this stage. Feature importance analysis identified three key parameters driving this discrimination (Fig S6A-C, Table 1). Based on these, we defined a composite metric, termed the cRhoA score, capturing membrane fluctuation amplitude, spatial organization of fluctuations, and oocyte shape anisotropy. By construction, positive cRhoA scores indicate increasing similarity to the phenotype of cRhoA oocytes, whereas negative scores indicate similarity to control oocytes. When computed at PBE-40 min, the cRhoA score efficiently separated control and cRhoA oocytes (Fig 5C), establishing it as a quantitative descriptor of the phenotype of cRhoA oocytes. Importantly, the cRhoA score correlated with the average velocity of cytoplasmic flows calculated from STICS analysis (Fig 2A-B), with higher scores associated with faster cytoplasmic flows (Fig S6D). This further supports the ability of the cRhoA score to capture the phenotype of cRhoA oocytes. We then used this score to probe unmanipulated oocytes. Applying the same pipeline to an independent dataset of unmanipulated mouse oocytes (Fig 5D), we found that while most displayed negative scores, a subset exhibited positive cRhoA scores (Fig 5E). These oocytes represented 11.5% of the population, indicating that the phenotype of cRhoA oocytes can arise spontaneously in mouse oocytes.

**Figure 5:**
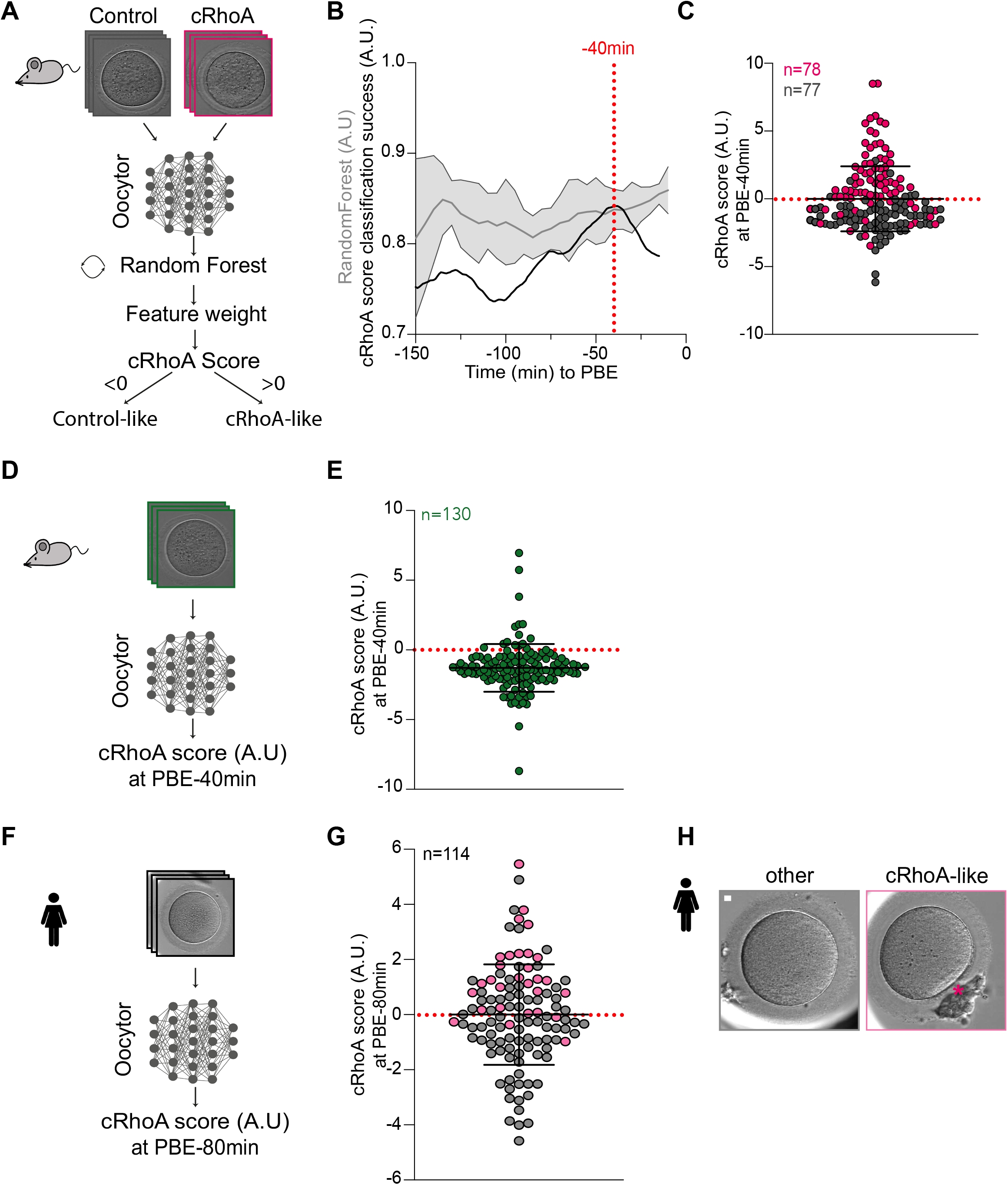
A subset of mouse and human oocytes naturally present a cRhoA phenotype. (A) Scheme of the Oocytor pipeline used to identify the most discriminating features between control (grey) and cRhoA (pink) mouse oocytes and compute a cRhoA score. (B) Evolution of the Random Forest classification performance (grey, A.U.; 5 repeats) and cRhoA score classification success (black, A.U.) between PBE-150 min and PBE-15 min. The Random Forest reflects classification performance between cRhoA and control oocytes, while the cRhoA score classification success indicates how well the reduced feature set discriminates between the two populations. The red dotted line marks the most discriminant time point (PBE-40 min). (C) cRhoA score at PBE-40 min for cRhoA (pink) and control (grey) oocytes. Data from 11 independent experiments. Mean ± SD is shown. The red dotted line represents a cRhoA score of 0. (D) Scheme of the Oocytor pipeline used to compute the cRhoA score at PBE-40 min for non-manipulated mouse oocytes (green). (E) cRhoA score at PBE-40 min for non-manipulated mouse oocytes (green). Films used for analysis were collected through the years (see Methods). Mean ± SD is shown. The red dotted line represents a cRhoA score of 0. 11.5% of oocytes display a positive score. (F) Scheme of the Oocytor pipeline used to compute the cRhoA score at PBE-80 min in human oocytes (black). (G) cRhoA score at PBE-80 min for human oocytes. Films used for analysis were collected from 2 separate datasets. Mean ± SD is shown. The red dotted line represents a cRhoA score of 0. 49% of oocytes display a positive score. Human oocytes are categorized as “cRhoA-like” (light pink) or “other” (grey). 88% of “cRhoA-like” human oocytes have a positive cRhoA score. (H) Representative images of human oocytes classified as “cRhoA-like” (light pink) and “other” (grey). The pink asterisk indicates a local flattening similar to the polarized domain observed in cRhoA mouse oocytes. Scale bar: 10 µm. For all graphs, n is the number of oocytes, each dot is 1 oocyte. PBE: Polar Body Extrusion.

**Table 1:** Features used by Oocytor for cRhoA oocyte discrimination. Values of mean morphological features ranked by feature importance used by Oocytor for cRhoA discrimination. The ones highlighted in yellow compose the cRhoA score.

**Figure 5**
| Feature Names | Feature Score |
| --- | --- |
| OoNormedFluctuationSpatialPower, | 39.3969089845981 |
| OoFluctuationFromLocalMean, | 30.3610611073633 |
| OoNormedAspectRatio, | 23.5593475665849 |
| OoPIVStrengthDirectionCenter, | 18.1986724389091 |
| OoFluctuationPropMoving, | 11.9843428265021 |
| OoRadiusMean, | 10.8570462392936 |
| OoPIVAngleEdge, | 10.1031326559933 |
| OoRadiusStd, | 9.79224323055017 |
| OoPIVStrengthDirection, | 9.42256608305216 |
| OoNormedPIVAngleEdge, | 9.27429062245294 |
| OoGLCMClusterShade, | 6.41615437745107 |
| OoCircularity, | 5.7223536551973 |
| OoNormMean, | 5.64209927085616 |
| OoMoranIndexK5, | 5.17695394540761 |
| OoPIVAngleCenter, | 4.98616815017982 |
| OoParticleAverageSize, | 4.8847721053482 |
| OoPIVMean, | 4.70525887124699 |
| OoNormedPIVDirCenter, | 4.67808205639431 |
| OoNormStd, | 4.30159525087682 |
| OoFluctuationPropGrowing, | 4.27589086112848 |
| OoCurvatureFlatProp, | 4.17592484519427 |
| PerivThicknessMin, | 4.05666285305155 |
| PerivThicknessCoefVar, | 4.048386262229 |
| OoNormCoefVar, | 3.96916273649895 |
| OoGLCMEntropy, | 3.96761584064064 |
| OoNormedFluctuationLocalMean, | 3.9236566503509 |
| OoParticleDistanceCoefVar, | 3.87009672490654 |
| PerivOoAreaRatio, | 3.82349663688301 |
| ZPGLCMClusterShade, | 3.80289373504988 |
| PerivThicknessMax, | 3.63806588517237 |
| OoKurtosis, | 3.48380381393172 |
| OoPIVCoefVar, | 3.40631976980491 |
| OoCurvatureMin, | 3.32837892493222 |
| OoFluctuationSpatialCorrLength, | 3.32741778316408 |
| OoParticleNumber, | 3.16273326574314 |
| ZPGLCMCorrelation, | 3.07451044694386 |
| OoMoranIndexK20, | 3.05505745380513 |
| OoPIVAverageDirectionCenter, | 3.05456992710924 |
| OoPIVAverageDirectionEdge, | 3.01964232102881 |
| ZPGLCMClusterTendency, | 2.99464041322774 |
| OoPIVAverageDirection, | 2.88216906882731 |
| OoCurvatureMax, | 2.83931319517993 |
| OoPIVStrengthDirectionEdge, | 2.75777372535556 |
| ZPGLCMContrast, | 2.62638307811702 |
| OoFluctuationSpatialMode, | 0.970523746037025 |

We next asked whether the cRhoA score could identify similar phenotypes in human oocytes. We reused the videos of maturing human oocytes from a previous study^23^ and added additional videos acquired under the same conditions at the same fertility center for a total number of 114 human oocytes. During fertility treatments, women undergo hormonal ovarian stimulation to promote oocyte development. After retrieval, not all oocytes are arrested in meiosis II and thus suitable for fertilization. The ones collected in Prophase I are considered to be of low quality and are not used for IVF. However, they can be used for research and are the only type of oocytes allowing the study of meiosis I from Prophase I onward in humans. We applied our pipeline to human oocytes undergoing meiosis I and computed a cRhoA score for each oocyte at PBE-80 min, corresponding to the same relative stage as PBE-40 min in mouse oocytes, since human meiosis I lasts approximately twice as long. This revealed a broad distribution of scores, with a substantial fraction of oocytes displaying positive values (49%, Fig 5G). To determine whether these scores correspond to visually identifiable phenotypes, we independently performed a blind manual annotation of all movies. Oocytes displaying characteristic features previously identified in cRhoA mouse oocytes, including cortical flattening and/or cortical travelling bands, were classified as “cRhoA-like”, whereas all others were classified as “other” (Fig 5H). With this visual phenotyping, we found that 34% of human oocytes were “cRhoA-like” (Fig 5G, pink dots). Importantly, oocytes classified as “cRhoA-like” by eye were strongly enriched in positive cRhoA scores (Fig 5G pink dots, 86% of “cRhoA-like” oocytes have a positive cRhoA score), whereas “other” oocytes predominantly displayed negative scores (Fig 5G grey dots, 63% of “other” oocytes have a negative cRhoA score), indicating good agreement between automated scoring and visual classification. Consistently, the discriminating power of the cRhoA score between manually annotated phenotypes peaked at PBE-80 min (Fig S6E).

Together, these results indicate that the morphological signature characteristic of cRhoA mouse oocytes can arise spontaneously in both mouse and human oocytes.

## Discussion

Our study shows that increased cortical tension in mammalian oocytes induces a precocious cortical polarity that drives cytoplasmic reorganization through aberrant cytoplasmic flows. This was observed following two distinct perturbations, a constitutive RhoA activation at the cortex and loss of the Arp2/3 complex, suggesting that precocious ectopic polarization does not arise from a specific molecular pathway but rather from a common mechanical state controlled by cortical tension. Cortical polarization therefore couples the cortex to the cytoplasm, potentially altering fertilization and developmental potential.

Actomyosin cortices behave as active materials that can spontaneously undergo mechanical instabilities and symmetry breaking^50^. Experimental studies have shown that increasing actomyosin contractility, and the resulting rise in cortical tension, is sufficient to trigger the transition from a homogeneous to a polarized cortical state^51,63^. Consistent with this framework, polarization emerged from several small cortical zones that progressively merged into a single domain. Similar cortical-flow-driven symmetry-breaking mechanisms have been described during embryonic polarity establishment, where anisotropies in cortical tension generate polarized actomyosin flows^64,65^. More recently, local regionalization of actomyosin organization has also been shown to generate spatial asymmetries during morphogenesis^54,66^. Together, these observations support the idea that precocious oocyte polarization results from a self-organized contractility-driven mechanism. Cortical instabilities may initiate symmetry breaking, whereas local mechanical asymmetries and mechanochemical feedback mechanisms^67,68^ could bias and stabilize the polarized domain.

We show here that ectopic cortical polarization generates cytoplasmic flows that redistribute the intracellular material over tens of micrometers. Oil droplets, organelles and meiotic spindles displayed similar transport dynamics, indicating that the process is largely object-independent and driven by physical advection. Such intracellular consequences may have been underestimated in other cellular contexts where cortical polarity has mainly been studied in relation to cell migration. Recent work in somatic cells similarly showed that actomyosin structures can generate non-specific cytoplasmic flows capable of redistributing soluble components through advection^69^. Force inference and hydrodynamic modeling showed that localized cortical forces are sufficient to generate cell-wide cytoplasmic streaming in mouse oocytes^70^, where actomyosin-dependent symmetry breaking contributes to meiotic spindle positioning, maintenance, and rotation^46,52,71^. Cytoplasmic flows can also organize intracellular components beyond the spindle, including mitochondria^23,72^. Our hydrodynamic model provides a mechanistic framework linking cortical polarization to cytoplasmic reorganization. Based on a coupled surface-bulk description of cortex and cytoplasm^56^, it recapitulates cortical deformations, cytoplasmic flows and spindle migration defects, demonstrating that mechanical coupling between a polarized contractile cortex and the viscous cytoplasm can drive large-scale intracellular reorganization. Together, these studies and our results support a broader view of cortical contractility as a general mechanical driver of intracellular organization in large mammalian oocytes.

The large-scale rearrangement of the oocyte interior persisted long after the initial symmetry-breaking event. In meiosis II, polarized oocytes displayed substantial organelle depletion, suggesting that cytoplasmic flows generated during meiosis I may alter intracellular organization through abnormal mechanical stress and/or directional transport toward the cortex, increasing the fraction of organelles lost during polar body extrusion. Whether this depletion is sufficient to impair developmental potential remains unclear, as polarized oocytes retained substantial numbers of organelles. However, reduced mitochondrial DNA content has previously been associated with impaired embryonic development in mammalian oocytes^73–77^. Furthermore, additional mechanisms resulting from increased cortical tension may also contribute to the lower quality of polarized oocytes.

Unexpectedly, polarized oocytes exhibited increased fertilization rates despite reduced developmental potential. In meiosis II, cRhoA remained polarized and was associated with asymmetric localization of Juno, CD9 and cortical granules^28,29,78^. Such enrichment could locally increase the probability of successful sperm-oocyte interactions, explaining the elevated fertilization rate. Consistent with this possibility, RhoA has been reported to accumulate at sites of sperm attachment and fusion, where it may help establish a membrane environment permissive for gamete fusion^46^. In addition, polarized oocytes displayed increased polyspermy. Because *in vitro* fertilization experiments were performed on zona-free oocytes, this phenotype is unlikely to result from defects in the cortical granule-dependent zona block to polyspermy, which acts tens of minutes after fertilization through zona pellucida hardening, and instead suggests an alteration of the fusion block, a membrane-based mechanism that limits additional sperm fusion within minutes after fertilization^79,80^. Nevertheless, defects in the zona block cannot be excluded. Notably, *arpc4-/-*oocytes also display abnormal cortical granule distribution in late meiosis I and exhibit defective polyspermy prevention *in vivo*^23^, raising the possibility that cortical polarization may affect multiple layers of polyspermy prevention. Together, these findings suggest that cortical organization contributes directly to fertilization success, and that perturbations of cortical organization may affect multiple aspects of fertilization.

A central implication of our findings is that mammalian oocytes, which typically maintain a homogeneous state throughout most of meiosis I, can transition into a polarized regime when cortical tension is elevated. In contrast to many non-mammalian oocytes that are polarized in Prophase and exhibit an anisotropic organization of their cytoplasm and cortex, mammalian oocytes lack such early polarity^12^. Our results provide a potential explanation for this absence of early polarity in mammals: maintaining a homogeneous cortical state may be crucial for preserving oocyte homeostasis prior to embryonic development. Although cortical polarization does not alter the geometry of the asymmetric meiotic division, it leads to cytoplasmic reorganization and perturbs fertilization outcomes. Thus, preventing precocious symmetry breaking may therefore be important for ensuring proper developmental potential. This interpretation aligns with the mammalian developmental strategy, in which major embryonic asymmetries arise after fertilization rather than being inherited from a pre-patterned oocyte^81^.

Finally, the observation that the morphological signature of polarized oocytes can arise spontaneously in both mouse and human oocytes highlights the potential physiological relevance of this mechanical state. The fact that ectopic polarization can be induced through two distinct perturbations suggests that multiple molecular pathways may converge on a common mechanical outcome. Spontaneously polarized oocytes could therefore arise from diverse defects affecting cortical contractility, actin organization or mechanochemical feedback mechanisms^35^. The relatively high incidence observed in human oocytes may reflect the heterogeneous quality of oocytes available for research and the diversity of cellular states present in these samples. Determining the molecular origins of this phenotype and its prevalence *in vivo* will be important for understanding whether precocious cortical polarization represents a previously unrecognized determinant of oocyte competence.

## Methods

### Mouse oocyte collection and culture

All animal studies were performed in accordance with the guidelines of the European Community and were approved by the French Ministry of Agriculture (authorization D750512). Ovaries were collected from 8-12 weeks old OF1 (Charles River Laboratories) and CD1 (Janvier-Labs) female mice and 8-16 weeks old control (wt/flox Cre-, wt/flox Cre +, flox/flox Cre-, wt/wt Cre+, or wt/wt Cre-) and conditional knockout for ArpC4 (flox/flox Cre+) females. The Cre recombinase was expressed only in growing oocytes thanks to the ZP3 promoter. Oocytes were extracted by dissecting the ovaries in homemade M2 medium^17^ supplemented with 1 µM Milrinone^82^ to synchronize them in Prophase I. Transferring the oocytes into a milrinone-free homemade M2 medium allowed meiosis resumption. All live culture and imaging were carried out in small droplets of medium covered with mineral oil at 37°C.

### *In vitro* fertilization (IVF) of *in vitro* matured oocytes

On day 1, Prophase I oocytes microinjected or not with cRNAs were incubated overnight in the incubator (37°C, 5% CO2) into a milrinone-free homemade M2 medium in order to progress to Metaphase II. Ferticult medium + 3% BSA was prepared on day 1 by supplementing Ferticult (JCD laboratoires; MT246-5) with BSA (A3311-10G; SIGMA), filtered with 0.22 µm filters (Clearline; 051732), and placed in the incubator overnight (at 37°C and 5% CO2), with the lid slightly open. On day 2 morning, the zona pellucida of the oocytes arrested in Metaphase II was removed by doing 3 rapid washes in Tyrode’s solution (Sigma-Aldrich; T1788). In parallel, a male mouse was sacrificed to collect the cauda epididymis with a piece of corpus epididymis and a piece of vas deferens, placed in a drop of 500 µL Ferticult + 3% BSA covered with oil at 37°C. Using two plastic cones to press on the epididymis, the sperm was released into the medium, and left to capacitate for 1h. Because zona-free oocytes generally display high fertilization rates and increased susceptibility to polyspermy, fertilization conditions were adjusted to yield relatively low fertilization rates and moderate polyspermy in control oocytes, thereby maximizing the sensitivity of the assay to detect differences between experimental groups. Therefore, for *in vitro* fertilization, the denuded oocytes were placed in a 100 µL drop of Ferticult + 3% BSA and then 0.2 µL of capacitated sperm was added. IVF proceeded for 1h in the incubator (37°C, 5% CO2). Afterwards, oocytes were washed in 5 separate drops of homemade M2 medium to remove any free sperm and then the cells were transferred into small drops of homemade M2 medium under mineral oil (M8410, Sigma) in a 35 mm tissue culture dish with cover glass bottom (FluoroDish FD35–100; World Precision Instruments), placing four oocytes per drop at opposite ends.

### Human oocyte collection, processing, culture and imaging

In this paper, we reused the videos of maturing human oocytes from^23^ and added more videos taken in the same conditions by the same fertility center. Human oocytes were collected during assisted reproduction treatment in the Centre Assistance Médicale à la Procréation Nataliance (Groupe Mlab, Pôle Santé Oréliance, Saran, France). Human oocytes not arrested in meiosis II and therefore not usable for patients were donated to research after informed consent. The videos of maturing human oocytes were anonymized and not related to the patient’s data. The oocytes were discarded after the movies.

Prophase I human oocytes were obtained from the hyaluronidase-dependent denudation of cumulus-oocyte complexes (COCs) harvested after ovarian stimulation (for details see^23^). Denuded oocytes were then rinsed in multiple drops of Sequential Fert and transferred to the intracytoplasmic sperm injection (ICSI) dish where they were exposed to a MOPS buffered media at 36.5°C during the time of the ICSI (G-MOPS Plus, Vitrolife, Sweden). All Metaphase I and Metaphase II oocytes were injected during the ICSI process and all oocytes including Prophase I were then rinsed in 4 microdrops of continuous embryo culture media (Sage One Step, Origio) before being transferred in the final incubation dish (Embryoslide Plus, Vitrolife, Sweden), containing Sage One Step under oil (Ovoil, Vitrolife, Sweden) pre-equilibrated overnight in the Embryoscope Plus Time-Lapse incubator (Vitrolife, Sweden) at 37°C, 6% CO2, 5% O2, 89% N2. Undisturbed continuous culture was then performed in the same incubation conditions for a maximum of 8 days (Day 0 to Day 7).

Videos of maturing Prophase I oocytes were collected retrospectively using the Embryo Viewer software (Vitrolife, Sweden), after an initial adjustment of the video focal plane on the Prophase I oocyte. Using the Embryoscope Plus incubator, images were acquired in 2048 x 1088 pixels (2.2 MP) with a 12-bit monochrome CMOS camera. A custom-made, high-quality 16x, 0.50 N.A. LWD Hoffman Modulation Contrast objective providing a resolution of 3 pixels per µm was used in conjunction with a single red LED for illumination (627 nm, duration 0.02 seconds per image). For each oocyte, the total light exposure time was < 40 seconds per day with a 10 min cycle time between image acquisitions. We kept only the videos where polar body extrusion occurred in the plane of observation.

### Constructs

The following constructs were used: pRN3-EzTD-mCherry-RhoA (cRhoA)^24,46^, pRN3-EzTD-mCherry (control)^22,47^, pspe3-GFP-UtrCH^18^, pGMHE-eGFP-Rab27a^23,60^, pRN3-H2B-GFP^83^.

### *In vitro* transcription of cRNAs and microinjection of cRNAs and oil droplets

Plasmids were linearized with SfiI or AscI restriction enzymes (NEBR0123S; NEBR0558S), and transcribed *in vitro* using the mMessage mMachine® T3 or T7 transcription kits (Thermo Fisher Scientific, AM1348; AM1344), allowing production of cRNAs. If needed, cRNAs were polyadenylated using a Poly(A) tailing kit (Thermo Fisher Scientific, AM1350). cRNAs were purified using the RNeasy® Mini kit (Qiagen, 74104) following the manufacturer’s instructions, eluted in 30 μL of RNase free water, aliquoted by 4 μL and stored at -20°C. Their quality was verified by electrophoresis gel migration and their concentration was quantified by NanoDrop 2000 (ThermoScientific). Final concentrations of cRNAs were >800 ng/µL, except for H2B-GFP (injected at 100 ng/µL).

cRNAs were centrifuged at 4°C at 13 000 rpm for at least 45 min prior to microinjection into the cytoplasm of oocytes blocked in Prophase I in homemade M2 medium supplemented with 1 µM of milrinone at 37°C under mineral oil. Microinjections were performed using an Eppendorf Femtojet microinjector^17^. Translation of cRNAs in oocytes was allowed for 2h to 4h, and then oocytes were transferred into a milrinone-free homemade M2 medium to allow meiosis resumption.

Oil droplet microinjection was performed as previously described^55^. Oocytes were first injected with cRNAs, then microinjected 2 h later with oil (Sigma, Fluorinert FC-70, F9880) presenting a density of 1.9x that of water at the maximal pressure of the microinjector.

### Live dyes and antibodies

To visualize the DNA, oocytes were incubated with SPY-DNA 1:2000 (Spirochrome, SC501); to visualize the microtubules, with SPY-Tubulin 1:1000 (Spirochrome, SC203) or 100 nM SiR-Tubulin (Spirochrome, SC002); to visualize the mitochondria, with 100 nM MitoTracker^TM^DeepRed FM (Thermo Fisher Scientific, M22426); to visualize the Golgi, with 500 nM NBD C6-Ceramide Complexed to BSA (Thermo Fischer Scientific, N22651); to visualize cortical actin, with SPY650-FastAct_X 1:1000 (Spirochrome; SC502). Oocytes were incubated for 30 min at 37°C in a medium containing the dyes, and then imaged in a medium containing the dyes, except for MitoTracker^TM^DeepRed FM.

To visualize CD9 and Juno in live, first the zona pellucida of the oocytes was removed by doing 3 rapid washes in Tyrode’s solution (Sigma-Aldrich; T1788). Then the denuded oocytes were incubated with 0.5 µg/µL AlexaFluor®647 α-mouse FR4 monoclonal antibody (Biolegend; 125009) to stain for Juno and 0.5 µg/µL of FITC α-mouse CD9 monoclonal antibody (Biolegend; 124807) to stain for CD9 for 15 min before acquisition^30,84^. The oocytes were fully imaged with 90 Z planes separated by 1 µm.

### Live imaging

Spinning disk microscope imaging was performed as previously described^16^. Briefly, oocytes were placed in a Fluorodish (World Precision Instruments #FD35-100) in small drops of homemade M2 medium covered with mineral oil. Images were acquired on 1/ a confocal spinning disk microscope (Leica DMI6000B) enclosed in a thermostatic chamber using a Plan-APO 40x/1.25-numerical aperture objective equipped with a Retiga 3 CCD camera (QImaging) coupled to a Sutter filter wheel (Roper Scientific) and a Yokogawa CSU-X1 Spinning disk or 2/ a confocal spinning disk microscope (Zeiss Axioobserver Z1) enclosed in a thermostatic chamber using a Plan-APO 40x/1.4-numerical aperture Plan-Apochromat objective equipped with a sCMOS Hamamatsu camera coupled to a Yokogawa CSU-W1 Spinning disk.

For meiotic maturation, films were acquired as follows: 1/ 4 Z planes separated by 4 µm every 30 min for 4h from nuclear envelope breakdown (BD) until BD+4h in transmitted light and fluorescence to monitor cRhoA polarization, then 1 Z plane every 2 min for 14h in transmitted light; 2/ 1 Z plane every 1 or 2 min for 20h starting from nuclear envelope breakdown (BD) in transmitted light; 3/ 8 or 10 Z planes separated by 4 µm every 10, 20 or 30 min for 20h starting from nuclear envelope breakdown (BD) in transmitted light and fluorescence. Only the oocytes that did nuclear envelope breakdown (BD) under 1h30 after milrinone washout were analyzed. Metamorph Software (Universal Imaging) was used to collect data.

### Immunofluorescent staining

Oocytes were washed in homemade M2 medium supplemented with Polyvinylpyrrolidone (PVP, Sigma, P0930) and adhered on coverslips coated with gelatine and polylysine^85^. The oocytes were fixed and permeabilized in 1x PBS, 100 mM Hepes pH 7, 50 mM EGTA pH 7, 10 mM MgSO4 buffer with 0.3% Triton X-100 and 2% Formaldehyde (Schuh 2008) for 30-60 min at 37°C. Oocytes were then washed 3 times for 10 min with 1x PBS - 0.1% Triton X-100, and further permeabilized with 1x PBS - 0.5% Triton X-100 for 10 min. Primary antibody staining was done in PBS - 0.1% Triton X-100, 3% BSA and left overnight at 4°C. The next day, oocytes were washed 3 times for 10 min with PBS - 0.1% Triton X-100, and secondary antibody staining was done in PBS - 0.1% Triton X-100, 3% BSA for 1h at room temperature in the dark. Before mounting, the coverslips were washed in 1x PBS for 10 min. Coverslips were mounted on glass slides with chambers (Electron Microscopy Sciences Slide Wells, 0.25mm Thick, N°70366-12) to avoid oocyte squashing. The oocytes were mounted with Prolong^TM^ Gold antifade reagent containing DAPI to stain for DNA (Thermo Fischer Scientific, P36931), left to polymerize overnight at 4°C, and sealed with nail polish.

Primary antibodies used were rabbit anti-Phospho-MLC2 (ser19) (1:200; Cell Signaling 3671). Secondary antibodies used were goat anti rabbit-Alexa 488 Highly Cross-Absorbed (1:500; Thermo Fischer Scientific A32731). Alexa-Fluor Plus 647 Phalloidin (1:50; Thermo Fischer Scientific A30107) was used to stain F-actin.

### Cortical tension measurements

Cortical tension at BD+2h was measured as previously described^20,22^. Briefly, after removing of the zona pellucida, oocytes were subjected to controlled micropipette aspiration using a glass micropipette connected to a pressure control system. The length of the aspirated cell protrusion was monitored as a function of the applied pressure, and the critical aspiration pressure was determined by extrapolation of the protrusion velocity to zero. Cortical tension was then calculated using the viscous drop model.

### Image analysis and quantifications

#### Aspect ratio measurement using Oocytor

The Aspect ratio of oocytes was measured using our open-source Fiji Plugin Oocytor^61^ (available in the GitHub repository: ). Using the “Get cortex” feature of Oocytor, we were able to extract the aspect ratio (AR) of the oocyte at every time point of transmitted light films, on a single Z plane. When AR equals 1, the shape is a circle. When AR is higher than 1, the shape is closer to an oval.

#### Measurement of polar body surface area

Measurements of polar body surface area were done on transmitted light films, 30 min after polar body extrusion (PBE), only on oocytes where the polar body was parallel to the plane of observation. Polar body surface area was measured by drawing an elliptical shape with the Fiji software^86^ and computing the area.

#### Measurement of oil droplet displacement and spindle migration duration

Oil droplet displacement duration was calculated by counting the number of frames during which the oil droplet visibly moved relative to the center of the oocyte using Fiji.

Spindle migration duration was calculated from films with oocytes incubated with SPY-Tubulin and SPY-DNA. The duration was calculated by counting the number of frames between when the spindle was bipolar and started migrating and when it reached the cortex.

#### Characterization of spindle migration

Oocytes incubated with SPY-Tubulin and SPY-DNA were used for these measurements. For both measurements, we used the straight-line tool on Fiji, and the center of our reference frame was the pole of the spindle closest to the center of the oocyte. When using the straight-line tool on Fiji, the angle displayed is that between the “horizontal” and the drawn segment: 0° to 180° counterclockwise and 0° to -180° clockwise. Only the oocytes where the spindle was in the plane of observation were measured.

#### Difference in angle between the long spindle axis and the migration axis

When the spindle was bipolar, at BD+4h, we traced a first segment along the long axis of the spindle starting at the pole closest to the center of the oocyte and measured the angle (α1). We traced a second segment along the migration axis of the spindle between BD+4h and BD+8h starting at the pole closest to the center of the oocyte and measured the angle (α2). We then calculated the absolute value of α1 - α2 allowing us to obtain |Δ_1_°|.

#### Difference in angle between the long spindle axis at BD+4h and before anaphase

When the spindle was bipolar, at BD+4h, we traced a first segment along the long axis of the spindle starting at the pole closest to the center of the oocyte and measured the angle (α1). Just before anaphase, we traced a second segment along the long axis of the spindle starting at the pole closest to the center of the oocyte and measured the angle (α3). We then calculated the absolute value of α1 - α3 allowing us to obtain |Δ_2_°|.

#### Quantification of fertilization, polyspermy and progression to the 2-cell stage

To visualize the pronuclei (PN) in order to compute the fertilization rate (percentage of zygotes displaying at least two PN, including normally fertilized and polyspermic zygotes) and assess polyspermy (percentage of zygotes with two PN and percentage of zygotes with more than two PN), oocytes were microinjected in Prophase I with cRNAs encoding for H2B-GFP with cRhoA, or H2B-GFP with Ezrin or nuclease free-water for controls. Translation of cRNAs was allowed for 2h, and then oocytes were transferred into a milrinone-free homemade M2 medium to allow them to reach Metaphase II overnight in the incubator. The next morning, all oocytes presenting a polar body were *in vitro* fertilized (IVF). We imaged the cells 8h30 after IVF in transmitted light and fluorescence (25 Z planes spaced 4 µm apart) and counted the number of pronuclei.

To quantify progression to 2-cell stage, oocytes were microinjected in Prophase I with cRNAs encoding for cRhoA or Ezrin or nuclease free-water for controls. Translation of cRNAs was allowed for 2h, and then oocytes were transferred into a milrinone-free homemade M2 medium to allow them to reach Metaphase II overnight in the incubator. The next morning, all oocytes presenting a polar body were *in vitro* fertilized (IVF) and placed in the incubator to develop. The next morning, approximately 24h after IVF, we counted the number of embryos at the 2- cell stage.

#### Quantification of active myosin-II in fixed oocytes

The background noise was manually subtracted from all immunofluorescence images. p-MLC2 fluorescence intensity at the cortex was measured in six random areas of the cortex using the same region of interest (ROI) of 4.35 µm^2^. Measurements were taken on one Z plane corresponding to the oocyte’s longest diameter. Using these six measurements, the average cortical p-MLC2 fluorescence intensity for each oocyte was calculated and normalized by the average of the controls for each coverslip.

#### Quantification of mitochondria and Golgi in live Metaphase II oocytes

The background noise was manually subtracted from all images. Oocytes in Metaphase II were incubated either in Mitotracker for 30 min at 37°C then washed out and imaged or incubated in the Golgi dye (NBD C6) for 15 min at 37°C and imaged with the dye still in the medium. To quantify total fluorescence intensities from the dyes, 66 Z planes spaced 1 µm apart and centered around the oocyte longest diameter were summed on a “SUM projection”. We only analyzed the oocytes with a polar body visible on the side of the oocyte (and not below or above it). An ROI encompassing the oocyte was drawn. The integrated density and ROI area were measured in order to calculate a ratio integrated density/area. Each oocyte ratio was normalized by the highest ratio of the experiment.

#### STICS analysis and calculation of average velocity

To quantify cytoplasmic movements by spatial temporal image correlation spectroscopy (STICS)^16,52^ we used the Fiji plugins downloaded from (http://research.stowers.org/imagejplugins)^87^ and implemented the appropriate ones for detecting cytoplasmic flows in mouse oocytes. We first aligned the transmitted-light movies of oocytes dividing in the plane of observation using the ‘‘Linear Stack Alignment with SIFT’’ plugin in Fiji and we subtracted the moving average using “stack subtract moving average jru v2” from the downloaded Fiji plugins. We then ran “stics map jru v2” on all films to be analyzed, after having selected a ROI corresponding to the cytoplasm of the oocyte for each video. This plugin measures the velocity vectors of the cytoplasm from temporal and spatial correlation throughout different timepoints, using a time correlation shift of 5. After running the plugin, we obtained STICS maps as well as an excel file with the numerical values of all the vectors. The STICS maps vectors were normalized in the settings on the plugin in order to obtain vectors of the same length and their velocity intensities were represented by a color gradient. In order to put all images on the same color scale, we applied the same contrast and brightness settings to all images. We then calculated the velocities of the vectors using the coordinates^87^ given by the plugin and calculated the average cytoplasmic movement. It allowed us to determine the highest one amongst our data set to establish the scale.

#### PIVlab analysis to quantify mitochondria movement

Particle Image Velocity (PIV) of mitochondria movement on 1 Z plane was measured using PIVlab application in MATLAB. Films were preprocessed by manually subtracting the background, removing the signal outside of the oocyte and orienting the films in order to have polar body extrusion on top, using Fiji. In image settings, we selected the “edit mask” mode and “expert” capabilities and used the dark area mask generator with a threshold of 0.01 to create a mask encircling only the oocyte’s cytoplasm for each frame. Then, using the PIV algorithm “multipass FFT window deformation” and creating a 3-pass window with 200, 100 and 50 pixels respectively, we analyzed all the frames. From the results and using the temporal derive parameters, we created a frame with the mean between the first and last frames of interest. Using this mean frame, and drawing an area around the oocyte’s cytoplasm using the polygon tool to delimit the area to analyze, we extracted the magnitude m/s and then converted it in µm/min.

### Machine learning approach for mouse oocyte morphological characterization and cRhoA score

We adapted our machine learning pipeline Oocytor^61^, distributed as an open-source Fiji plugin (available in the GitHub repository: https://github.com/gletort/Oocytor) to automatically measure and compare morphological features of control (oocytes expressing Ezrin and non-injected oocytes) and cRhoA oocytes, as previously described^23,62^. Oocytor allowed us to detect the oocyte contours and extract features describing their morphology and dynamics^61^ at each time point. Movies were acquired in transmitted light every 2 min and were cropped between BD+4h and just before polar body extrusion (PBE). To enable comparison with dataset acquired over several years^61^, movies were temporally and spatially rescaled by selecting frames every 6 min and setting a spatial resolution of 0.227 µm/pixel. As we were interested in the ectopic polarization occurring in cRhoA oocytes opposite to where polar body extrusion takes place, we analyzed only the oocytes in which PBE occurred in the plane of observation (78 cRhoA films and 77 control films) and aligned all movies using PBE time as a temporal reference. To further enforce the measure of oocyte features relative to PBE, we introduced normalized features by dividing each feature value at all time points before PBE by its value immediately before PBE, such that all normalized features equal 1 at PBE^23,61^. Once we measured the morphological and dynamics features, and their corresponding normalized features, we trained a Random Forest classifier to distinguish control and cRhoA oocytes, as previously described^23,61,62^. Classification performance was evaluated with a 5-fold cross-validation scheme at multiple time points between PBE-150 min and PBE-10 min (Fig 5B). The most discriminant features were identified based on feature importance (Table 1; Fig S6A-C).

Based on the three most discriminant features, we defined a composite metric termed the cRhoA score: cRhoA score = OoFluctuationFromLocalMean - OoNormedFluctuationSpatialPower - OoNormedAspectRatio. OoFluctuationFromLocalMean reflects short-timescale membrane movements^23^. Short timescale fluctuations were higher in cRhoA oocytes around PBE-40 min. OoFluctuationSpatialPower captures the spatial scale of these fluctuations^23^. OoNormedFluctuationSpatialPower corresponds to its normalized value compared to PBE. The spatial fluctuations strongly increased between PBE-40 min and PBE in cRhoA oocytes, thus the normalized feature values were smaller. OoAspectRatio corresponds to the ratio between the major and minor axes of the ellipse fitted to the oocyte contour. OoNormedAspectRatio corresponds to its normalized value compared to PBE. The aspect ratio of control oocytes was stable, while cRhoA oocytes were more circular at PBE-40 min and strongly deformed around PBE. Thus, the normalized feature values were lower for cRhoA oocytes. Normalized features were computed as described above. To assess whether this phenotype occurs naturally, we computed the cRhoA score on an independent dataset of 130 non-injected, non-manipulated mouse oocytes^61^, processed identically (selection based on PBE plane, scaling, temporal alignment, and Oocytor analysis). The cRhoA score was computed at PBE-40 min, corresponding to the time point of maximal discrimination (Fig 5B, E).

### Machine learning approach for human oocyte analysis and cRhoA score

Human oocytes were imaged in transmitted light in a clinical context. Individual wells were automatically extracted from movies containing 1 to 4 wells. When multiple oocytes were present in a single well, they were separated using a Fiji macro that automatically detects individual oocytes, ensuring that each movie contained a single centered oocyte. Oocytes located at the border of the well were excluded. As for mouse oocytes, only oocytes with PBE in the plane of observation were analyzed (n=114 oocytes from two independent datasets). Movies were temporally aligned with respect to PBE, and only frames between NEBD+9h and PBE were retained, as earlier time points may include cortical contractions^23^. Movies had a temporal resolution of 10 min and a spatial resolution of 0.34375 µm/pixel. Movies were processed using the same Oocytor pipeline, and the same features were extracted. The cRhoA score was computed at PBE-80 min, twice that of mouse oocytes, since human meiosis I lasts approximately twice as long as mouse meiosis I. Movies were then manually annotated in a blinded manner and classified into two categories based on qualitative inspection: “cRhoA-like” or “other” (Fig 5H). The cRhoA scores were plotted, and color-coded according to these categories (Fig 5G). We then evaluated the discriminating power of the cRhoA score over time, showing that classification performance between these two categories indeed peaked at PBE-80 min (Fig S6E).

### Statistical analysis

Experiments (N) were repeated at least three times. All statistical analyses were performed using GraphPad Prism 10.1.0. Data normality was assessed using the Shapiro-Wilk test. For comparison between two independent groups, statistical significance was assessed using an unpaired t-test for normally distributed data or the nonparametric Mann-Whitney test for non-normally distributed data. For paired data, statistical significance was assessed using a paired t test for normally distributed data or a Wilcoxon matched-pairs signed rank test for non-normally distributed data. Correlations between variables were assessed using Spearman’s rank correlation coefficient (*r*s). Exact Cochran-Mantel-Haenszel tests stratified by experiment were used for contingency table analyses for N=3 experiments. Statistical significance was assessed using a significance level of α = 0.05. The number of oocytes analysed (n) is indicated in the figure legends. Error bars represent mean ± SD. A P-value < 0.05 was considered statistically significant. In all figures, *P < 0.05, **P < 0.01, ***P < 0.001, ****P < 0.0001; ns, not significant.

## Author Contributions

M-E.T conceived the project, which was supervised by M-E.T and M-H.V. A.S designed, performed and analyzed all experiments on mouse oocytes except for *arpc4-/-*oocytes that were investigated by E.Nik. G.L performed all analysis using the Oocytor pipeline. E.Nei and H.T designed the biophysical model. E.Nei performed all numerical simulations and subsequent analysis. S.M and H.T designed the optimal-transport method to measure cortical flows. M.Y and X.P-V provided the human oocyte transmitted light videos. M-E.T and A.S wrote the manuscript, which was seen and corrected by all authors.

## Acknowledgments

We thank Raphaël Voituriez, Guillaume Halet and all members of the Verlhac-Terret laboratory for discussions, Christine Gourier and Adrien Freyss for their advice on IVF experiments, Anne-Marie Berisha and Oriol Mañé Benach for preliminary experiments, the CIRB animal and Orion facilities, and especially Héloïse Monnet and Thomas Caille for advice on the PIVlab analysis to quantify mitochondria movement. The Verlhac-Terret and Turlier laboratories are supported by CNRS, INSERM, Collège de France and the Bettencourt Schueller Foundation. This work was supported by the French Government under the France 2030 Investment Plan, with funding managed by the French National Research Agency (ANR), through project ANR-24-PESF-0001 within the PEPR Women’s Health, Couples’ Health. The authors acknowledge support from the Major Research Program « DEVINE » launched by PSL Research University and implemented by ANR, with the reference ANR-10-IDEX-0001, and through grants from the ANR (ANR-18-CE13 to M-H.V; ANR-24-CE13-5691 to M-E.T), PSL-QLife (Q-life ANR-17-CONV-0005 to M-E.T) and FRM (DEQ201903007796 to M-H.V; SMC202405019631 to M-E.T). A.S was supported by fellowships from French Ministère de la Recherche and from ARC (ARCDOC42025120010988). E.Nei received funding from the European Union’s Horizon EU research and innovation program under grant agreement 101105565. H.T received funding from the European Research Council (ERC) under the European Union’s Horizon 2020 research and innovation program (grant agreement no. 949267).

## Competing interests

The authors declare no competing interests.

## Data and materials availability

All data are available in the main text or the supplementary materials.

## Figure, Table and Movie legends

**Figure S1:**
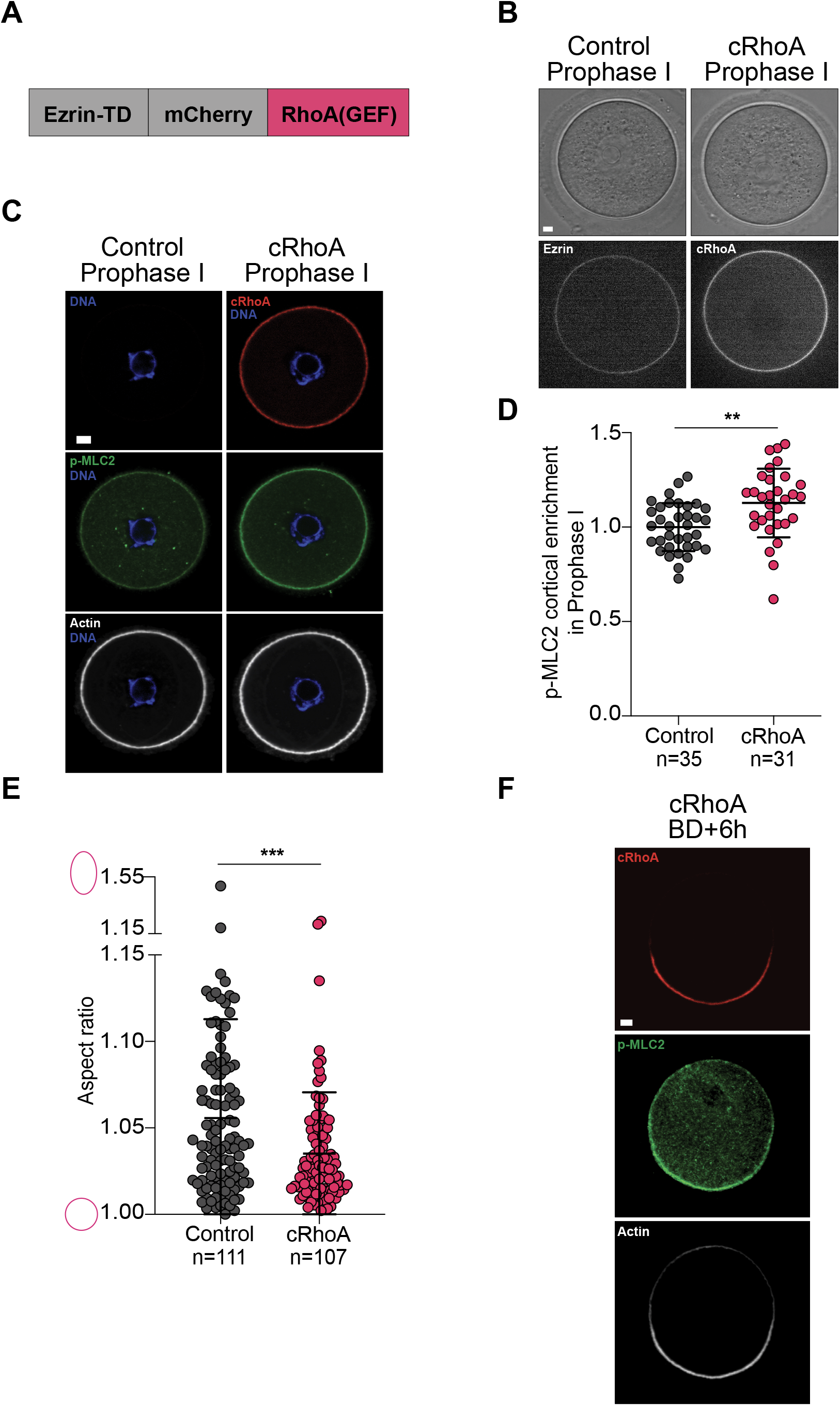
cRhoA localizes to the cortex in Prophase I, recruiting active myosin-II, and polarizes mid-meiosis I along with actin and active myosin-II. (A) Scheme of the cRhoA construct. (B) Transmitted light and fluorescent images of control and cRhoA oocytes in Prophase I. Control oocytes express Ezrin (white), cRhoA oocytes express cRhoA (white). One Z plane is shown. (C) Control and cRhoA oocytes fixed in Prophase I and stained for active myosin-II (p-MLC2, green), Actin (phalloidin, white) and DNA (DAPI, blue). cRhoA oocytes express cRhoA (red). One Z plane is shown, with the same contrast for all images. (D) Active myosin-II (p-MLC2) enrichment at the cortex in control and cRhoA oocytes in Prophase I. Data from 3 independent experiments. Mean ± SD is shown. Statistical significance of differences was assessed by an unpaired t test, P = 0.0014. (E) Aspect ratio of control and cRhoA oocytes in Prophase I, measured using Oocytor. Data from 10 independent experiments. Mean ± SD is shown. Statistical significance of differences was assessed by a Mann-Whitney test, P = 0.0005. (F) cRhoA oocyte expressing cRhoA (red) fixed at BD+6h and stained for active myosin-II (p-MLC2, green) and Actin (phalloidin, white). One Z plane is shown. For all images, scale bar: 10 µm. For all graphs, n is the number of oocytes, each dot is 1 oocyte. BD: Nuclear envelope breakdown.

**Figure S2:**
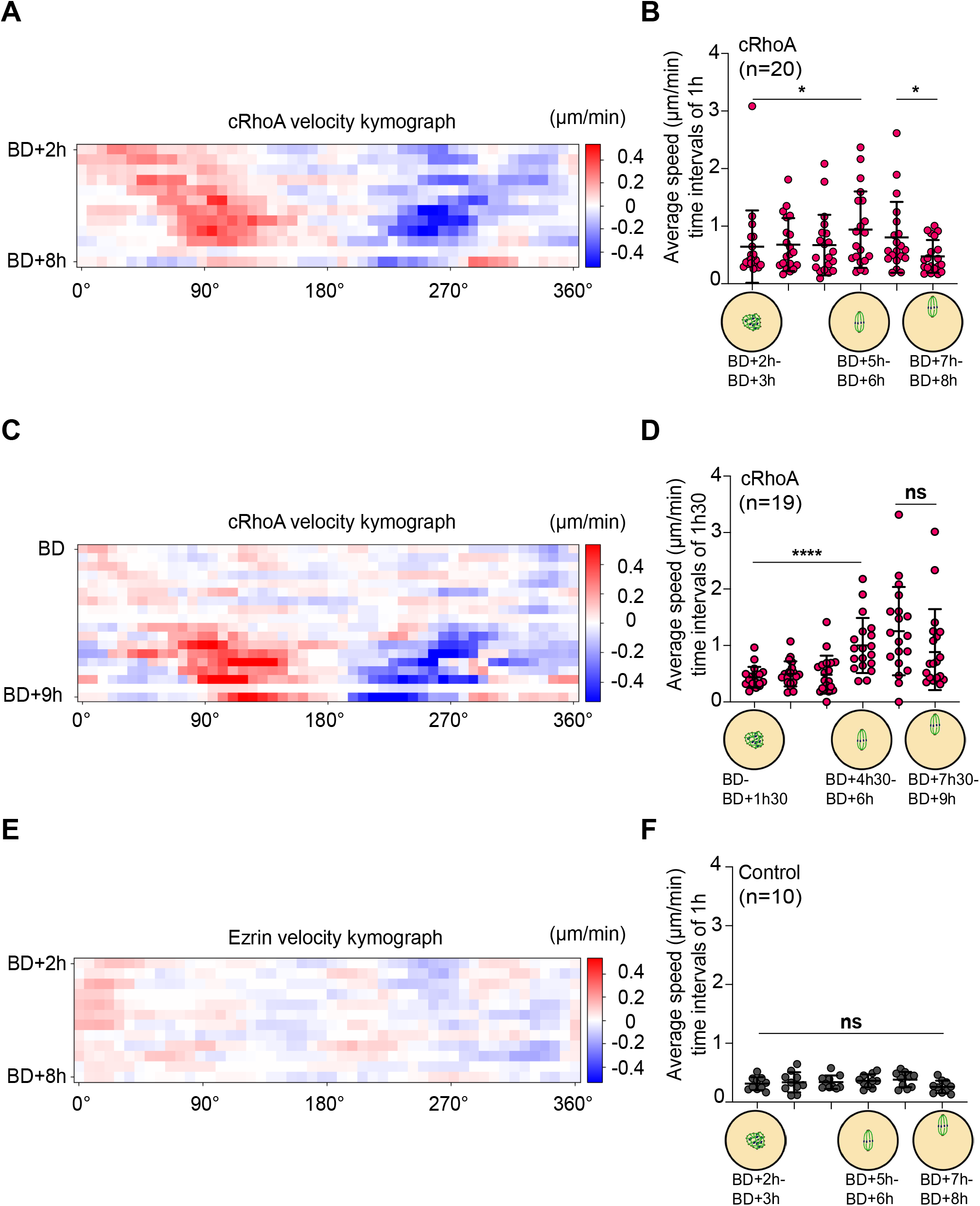
Control oocytes do not polarize in meiosis I whereas cRhoA oocytes analyzed at different time resolutions polarize precociously mid-meiosis I. (A) Averaged cRhoA velocity kymograph at the cortex between BD+2h and BD+8h, computed from 20 films with images acquired every 30 min, collected from 3 independent experiments. The X-axis axis represents the unfolded oocyte cortex as shown on the scheme Fig 1H left, and the Y-axis represents the time from BD+2h until BD+8h and the velocity of cRhoA fluorescence (µm/min). (B) Average speed of polarization of cRhoA at the cortex between BD+2h and BD+8h calculated from the averaged cRhoA velocity kymograph in A. Data from 20 films, collected from 3 independent experiments. Statistical significance of differences was assessed by a Wilcoxon matched-pairs signed rank test, from left to right P = 0.0362, P = 0.0107. (C) Averaged cRhoA velocity kymograph at the cortex, between BD and BD+9h, computed from 19 films with images acquired every 10 min, but analyzed as if taken every 30 min, collected from 3 independent experiments. The X-axis represents the unfolded oocyte cortex as shown on the scheme Fig 1H left, and the Y-axis represents the time from BD until BD+9h and the velocity of cRhoA fluorescence (µm/min). (D) Average speed of polarization of cRhoA at the cortex between BD and BD+9h calculated from the averaged cRhoA velocity kymograph in C. Data from 19 films, collected from 3 independent experiments. Statistical significance of differences was assessed by a Wilcoxon matched-pairs signed rank test, from left to right P < 0.0001, P = 0.0728. (E) Averaged Ezrin velocity kymograph at the cortex in control oocytes between BD+2h and BD+8h, computed from 10 films with images acquired every 30 min, collected from 3 independent experiments. The X-axis represents the unfolded oocyte cortex as shown on the scheme Fig 1H left, and the Y-axis represents the time from BD+2h until BD+8h and the velocity of Ezrin fluorescence (µm/min). (F) Average speed of polarization of Ezrin at the cortex between BD+2h and BD+8h calculated from the averaged Ezrin velocity kymograph in E. Data from 10 films, collected from 3 independent experiments. Statistical significance of differences was assessed by a paired t test, P = 0.1549. For all graphs, cRhoA oocytes are in pink, controls in grey, n is the number of oocytes, each dot is 1 oocyte. BD: Nuclear envelope breakdown; PBE: Polar Body Extrusion.

**Figure S3:**
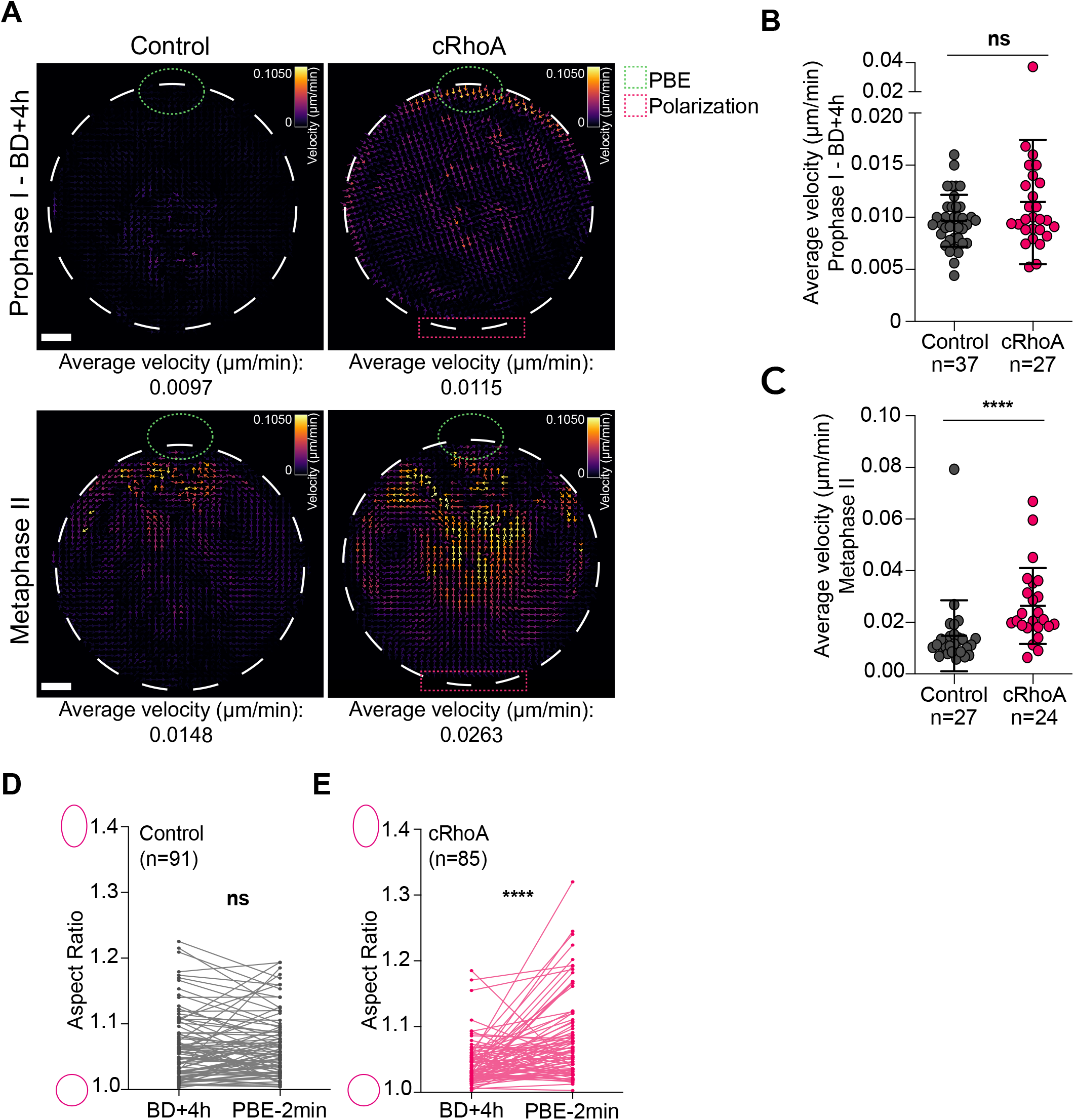
Cytoplasmic flows are comparable in control and cRhoA oocytes before mid-meiosis I, but stronger in Metaphase II for cRhoA oocytes; cRhoA oocytes change shape after ectopic polarization. (A) Representative vector maps of cytoplasmic flows from transmitted light videos obtained using STICS analysis for control and cRhoA oocytes, measured between Prophase I and BD+4h, and in Metaphase II during 3h. Scale bars: 10 µm. (B) Average velocity, calculated from all average velocity vectors between Prophase I and BD+4h for control and cRhoA oocytes. Data from 3 independent experiments. Means ± SD are shown. Statistical significance of differences was assessed by a Mann-Whitney test, P = 0.2861. (C) Average velocity, calculated from all average velocity in Metaphase II for control and cRhoA oocytes. Data from 6 independent experiments. Means ± SD are shown. Statistical significance of differences was assessed by a Mann-Whitney test, P < 0.0001. (D) Evolution of the Aspect ratio of control oocytes between BD+4h and PBE-2 min. Data from 14 independent experiments. Statistical significance of differences was assessed by a Wilcoxon matched-pairs signed rank test, P = 0.7296. (F) Evolution of the Aspect ratio of cRhoA oocytes between BD+4h and PBE-2 min. Data from 14 independent experiments. Statistical significance of differences was assessed by a Wilcoxon matched-pairs signed rank test, P < 0.0001. For all graphs, cRhoA oocytes are in pink, controls in grey, each dot is 1 oocyte, n is the number of oocytes. BD: Nuclear envelope breakdown; PBE: Polar Body Extrusion.

**Figure S4:**
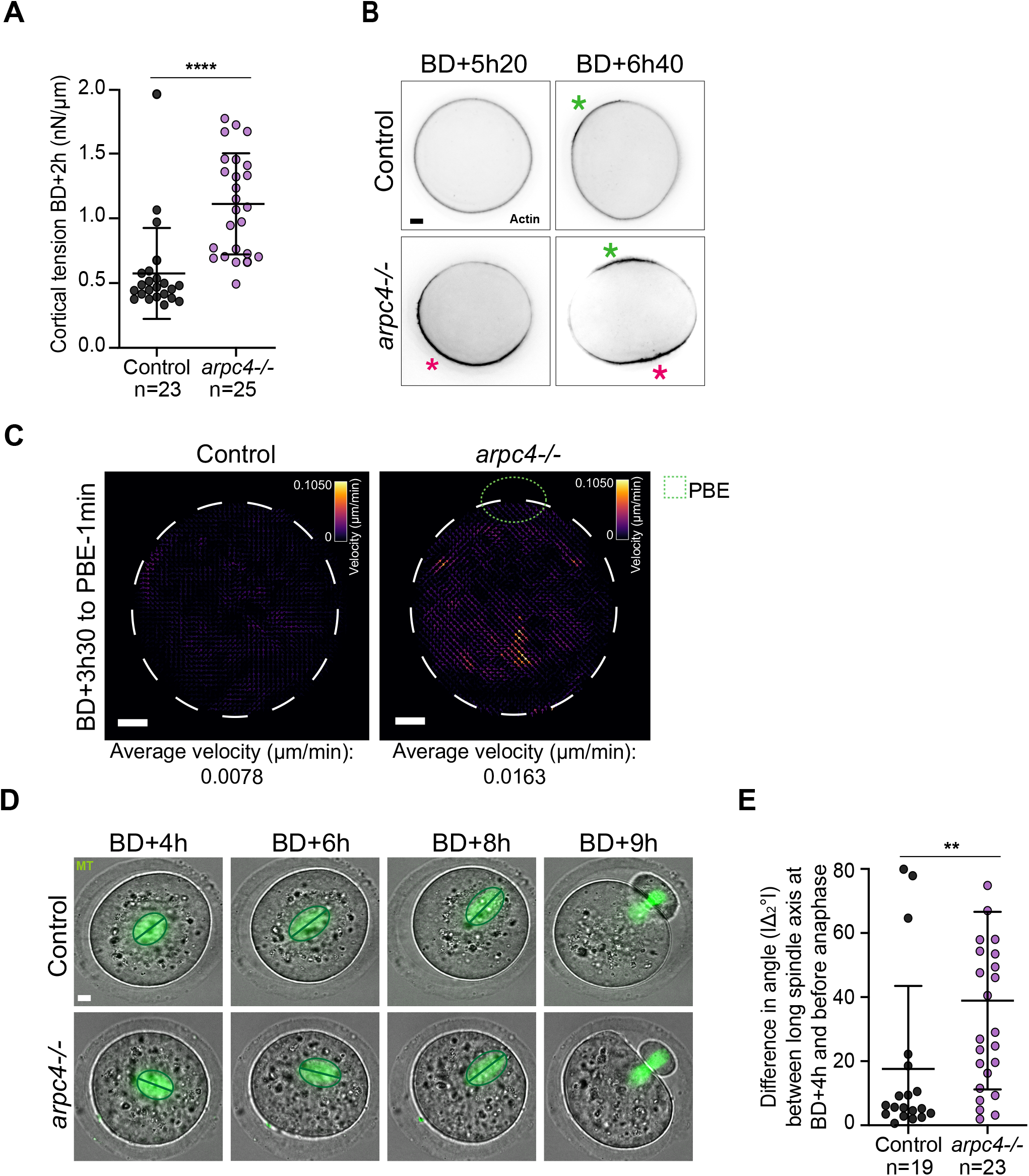
*arpc4-/-*oocytes, which have a higher cortical tension, behave as cRhoA oocytes. (A) Cortical tension (nN/μm) measured by micropipette aspiration for control and *arpc4-/-*oocytes. Data from 3 independent experiments. Means ± SD are shown, each dot is 1 oocyte. Statistical significance of differences was assessed by a Mann-Whitney test, P < 0.0001. (B) Fluorescent images of control and *arpc4-/-*oocytes at BD+5h20 and BD+6h40 expressing GFP-UtrCH to label actin filaments. Cortical actin polarizes in *arpc4-/-*oocytes (pink asterisk). The green asterisk indicates the actin cap forming when the chromosomes are in proximity with the cortex at the end of meiosis I. (C) Representative vector maps of cytoplasmic flows from transmitted light videos obtained using STICS analysis for control and *arpc4-/-*oocytes, measured between BD+3h30 and PBE-1 min. (D) Time-lapse fluorescent and corresponding transmitted light images of control and *arpc4-/-*oocytes between BD+4h and BD+9h. Microtubules (MT, green) were visualized with SiR-Tubulin. Images are 12 µm Z-projections. (E) Difference in angle (|Δ_2_°|) between the long axis of the spindle at BD+4h and just before anaphase, as computed in Fig 2H. Data from 5 independent experiments. Means ± SD are shown. Statistical significance of differences was assessed by a Mann-Whitney test, P = 0.0025. For all graphs, *arpc4-/-*oocytes are in purple, controls in grey, each dot is 1 oocyte, n is the number of oocytes. For all images, scale bar: 10 µm. BD or NEBD: Nuclear envelope breakdown; PBE: Polar Body Extrusion.

**Figure S5:**
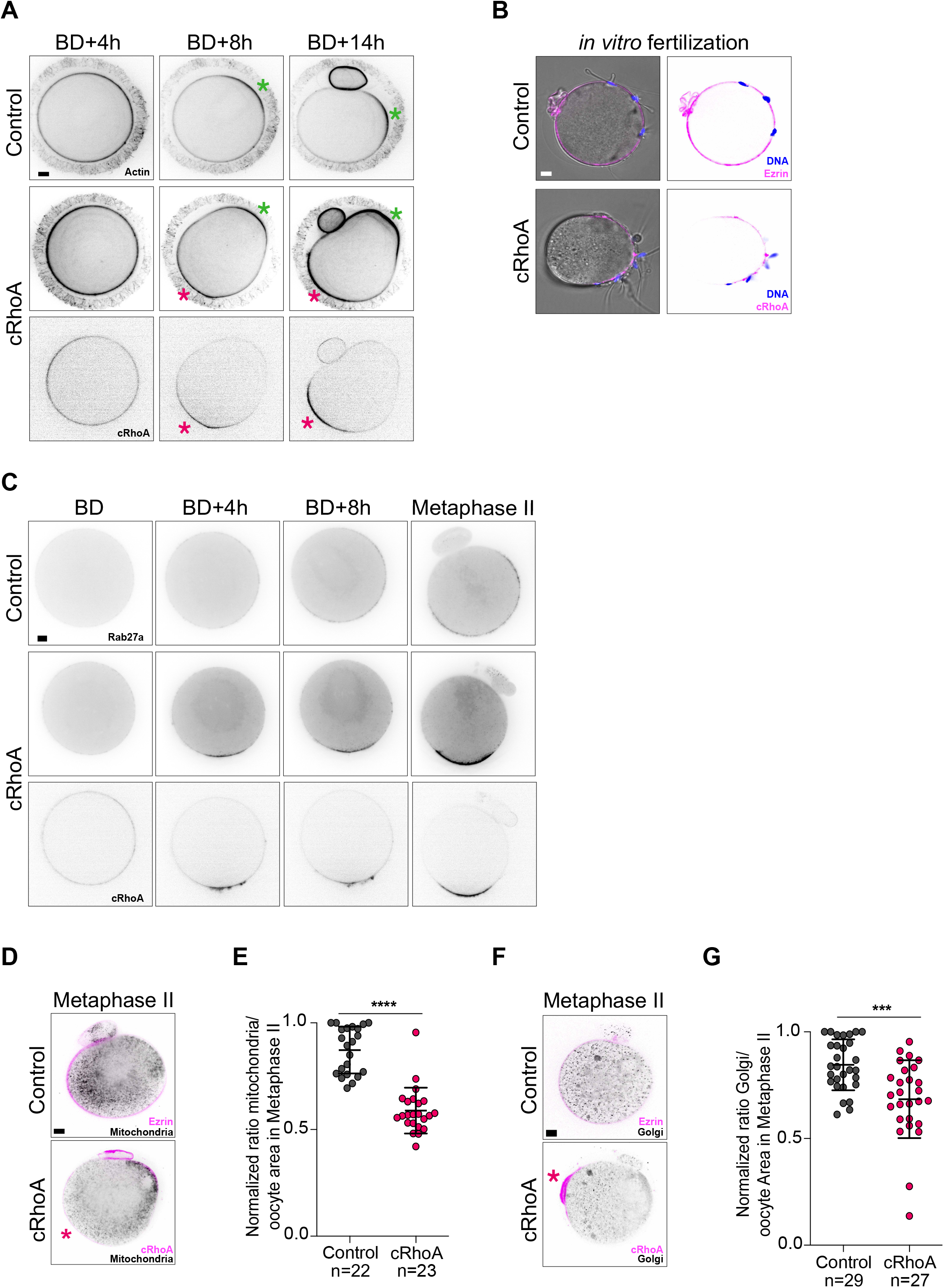
cRhoA oocytes form two actin caps in meiosis I and II, they are depleted in organelles in meiosis II, and sperm can bind to their polarized region in meiosis II. (A) Time-lapse fluorescent images of control and cRhoA oocytes between BD+4h and BD+14h. Cortical Actin was visualized with SPY-FastAct_X (black). cRhoA oocytes express cRhoA (black, bottom), which polarizes (pink asterisk). The green asterisk indicates the actin cap forming when the chromosomes are in proximity with the cortex at the end of meiosis I and in meiosis II. (B) Fluorescent images of control and cRhoA oocytes fertilized *in vitro* in Metaphase II. Control oocytes express Ezrin (pink), cRhoA oocytes express cRhoA (pink). DNA (blue) was visualized with SPY-DNA. One Z plane is shown. (C) Time-lapse fluorescent images of control and cRhoA oocytes between BD and Metaphase II. All oocytes express Rab27a (cortical granules, black), cRhoA oocytes express cRhoA (black). One Z plane is shown. (D) Fluorescent images of control and cRhoA oocytes in Metaphase II. Mitochondria (black) were visualized with Mitotracker. Control oocytes express Ezrin (pink), cRhoA oocytes express cRhoA (pink), which polarizes (pink asterisk). Images are 9 µm Z-projections. (E) Normalized ratio of mitochondria/oocyte area in control and cRhoA oocytes in Metaphase II. Data from 3 independent experiments. Mean ± SD is shown. Statistical significance of differences was assessed by a Mann-Whitney test, P < 0.0001. (F) Fluorescent images of control and cRhoA oocytes in Metaphase II. Golgi (black) was visualized with (NBD C6). Control oocytes express Ezrin (pink), cRhoA oocytes express cRhoA (pink), which polarizes (pink asterisk). Images are 9 µm Z-projections. (G) Normalized ratio of Golgi/oocyte area in control and cRhoA oocytes in Metaphase II. Data from 4 independent experiments. Mean ± SD is shown, each dot is 1 oocyte. Statistical significance of differences was assessed by a Mann-Whitney test, P = 0.0002. For all graphs, cRhoA oocytes are in pink, controls in grey, n is the number of oocytes. For all images, scale bar: 10 µm. BD: Nuclear envelope breakdown.

**Figure S6:**
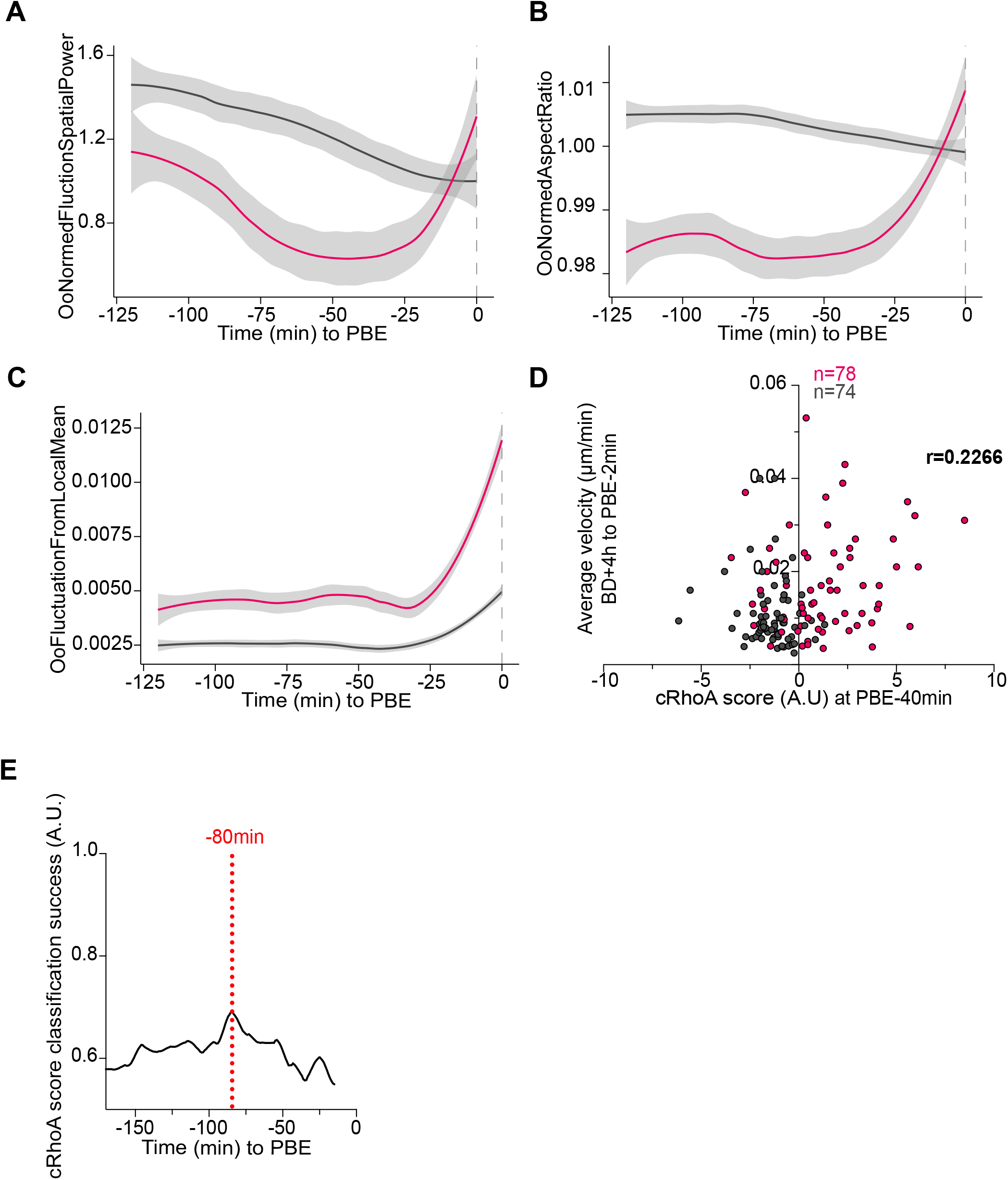
Evolution in time of the features composing the cRhoA score, correlation between cytoplasmic flows and cRhoA score in meiosis I and discriminating power of the cRhoA score in human oocytes. (A-C) Temporal evolution of the three features composing the cRhoA score between PBE-125 min and PBE in control (grey) and cRhoA (pink) oocytes. Mean plotted with confidence interval of 95%. (D) Average cytoplasmic velocity of control (grey) and cRhoA (pink) oocytes between BD+4h and PBE-2 min (from Fig 2B), plotted against their cRhoA score at PBE-40 min. Each dot is 1 oocyte, n is the number of oocytes. Spearman correlation calculated (r), P < 0.0075. (E) Evolution of cRhoA score classification success (black, A.U.) between “cRhoA-like” and “other” oocytes over time. The red dotted line marks the most discriminant time point (PBE-80 min).

**Movie 1: cRhoA oocytes divide normally.** Time-lapse transmitted light movie of control and cRhoA oocytes between Prophase I and BD+12h.

**Movie 2: cRhoA and *arpc4-/-*oocytes display aberrant cytoplasmic flows from mid-meiosis I on.** Representative transmitted light videos used for STICS analysis for control and cRhoA oocytes between Prophase I and BD+4h (first row), BD+4h and PBE-2 min (second row) and in Metaphase II during 3h (third row), and for control and *arpc4-/-*oocytes between BD+3h30 and PBE-1 min (fourth row).

**Movie 3: Oil droplets are pushed toward the future division site in cRhoA oocytes.** Time-lapse transmitted light movie of control and cRhoA oocytes injected with an oil droplet from BD+4h until BD+14h.

**Movie 4: Spindles are pushed toward the future division site in cRhoA oocytes.** Time-lapse fluorescent movie of control and cRhoA oocytes between BD+2h and BD+10h. Microtubules (MT, white) were visualized with SPY-Tubulin and DNA (green) was visualized with SPY-DNA. Control oocytes express Ezrin (white), cRhoA oocytes express cRhoA (white). Frames are 12 to 15 µm Z-projections.

**Movie 5: Spindle movement depends on its initial position in cRhoA oocytes.** Representative simulations illustrating spindle dynamics for different initial configurations, highlighting the dependence of rotational behavior on initial conditions.

**Movie 6: Mitochondria are pushed toward the future division site in cRhoA oocytes.** Time-lapse fluorescent movie of control and cRhoA oocytes between BD+1h and BD+8h. Mitochondria (black) were visualized with Mitotracker. Control oocytes express Ezrin (pink), cRhoA oocytes express cRhoA (pink). Frames are 8 to 12 µm Z-projections.

**Movie 7: cRhoA oocytes form two actin caps in meiosis I and II.** Time-lapse fluorescent movie of control and cRhoA oocytes between BD+4h and BD+14h. Cortical Actin was visualized with SPY-FastAct_X (black). For all movies, BD: Nuclear envelope breakdown, PBE: polar body extrusion.

## DETAILED METHODS FOR FLOW INFERENCE WITH OPTIMAL TRANSPORT AND A HYDRODYNAMIC MODEL LINKING CORTICAL POLARIZATION TO CYTOPLASMIC REORGANIZATION IN CRHOA OOCYTES

**RelateD to Fig. 1, Fig. S2 anD Fig. 3**

### Flow inference with optimal transport

#### 1. Pre-Processing

We analyzed 3D time-lapse videos of cRhoA oocyte development. The time-lapse recordings vary in their start times: some begin immediately after Nuclear Envelope Breakdown (NEBD), while others start later, such as two hours post-NEBD (NEBD + 2h). The videos continue up to ten or thirty minutes before polar body extrusion (PBE). We start with:

- **Cropping**: before the first division (PBE)
- **Rotating**: such as PBE happens at the top center (Supplementary Text Fig. 1).
- **Projection**: we projected the 3D time-lapses into 2D to simplify processing and avoid handling the oocytes as 3D objects. We specifically used projections onto two planes rather than a full Z-stack projection to minimize noise.

**Supplementary Text Figure 1.**
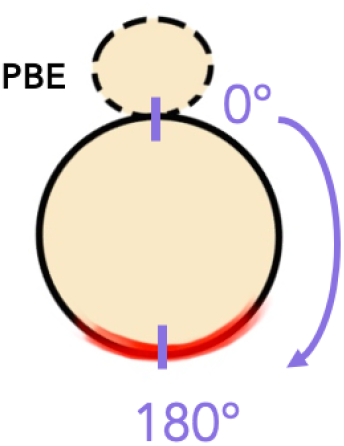
Orientation of the oocytes for our analysis tool (illustrated in Supplementary Text Fig. 2)

First, we segment the cortex (where polarization occurs) in each frame independently. To improve image quality, we apply:

- **CLAHE (Contrast Limited Adaptive Histogram Equalization)** [1]: Enhances local contrast by dividing the image into small patches and independently equalizing their histograms.
- **Non-Local Means (NLM) Denoising** [2]: Removes noise by averaging pixels across the entire image based on similarity of their surrounding patches (e.g., a 5 × 5 neighborhood).

Next, we use:

- **Otsu’s thresholding**[3]: Automatically selects a global threshold to separate the cortex (foreground) from the background by maximizing inter-class variance (i.e., making foreground/background pixel intensities as distinct as possible).
- **Contour detection** [4]: Converts the binary thresholded image into a continuous boundary (polygon) by tracing connected pixel edges.
- **Angular Sectors**: we construct a set of *n* = 50 regularly spaced angular sectors (angular increments of 7.2^◦^) and aggregate pixel intensity values within each sector to obtain an intensity distribution. As illustrated in Supplementary Text Fig. 2, these distributions enable frame-by-frame analysis of intensity variations. To track the motion of pixel intensities in consecutive frames, we employ computational optimal transport methods, allowing us to determine the displacement of intensity values over time.

**Supplementary Text Figure 2.**
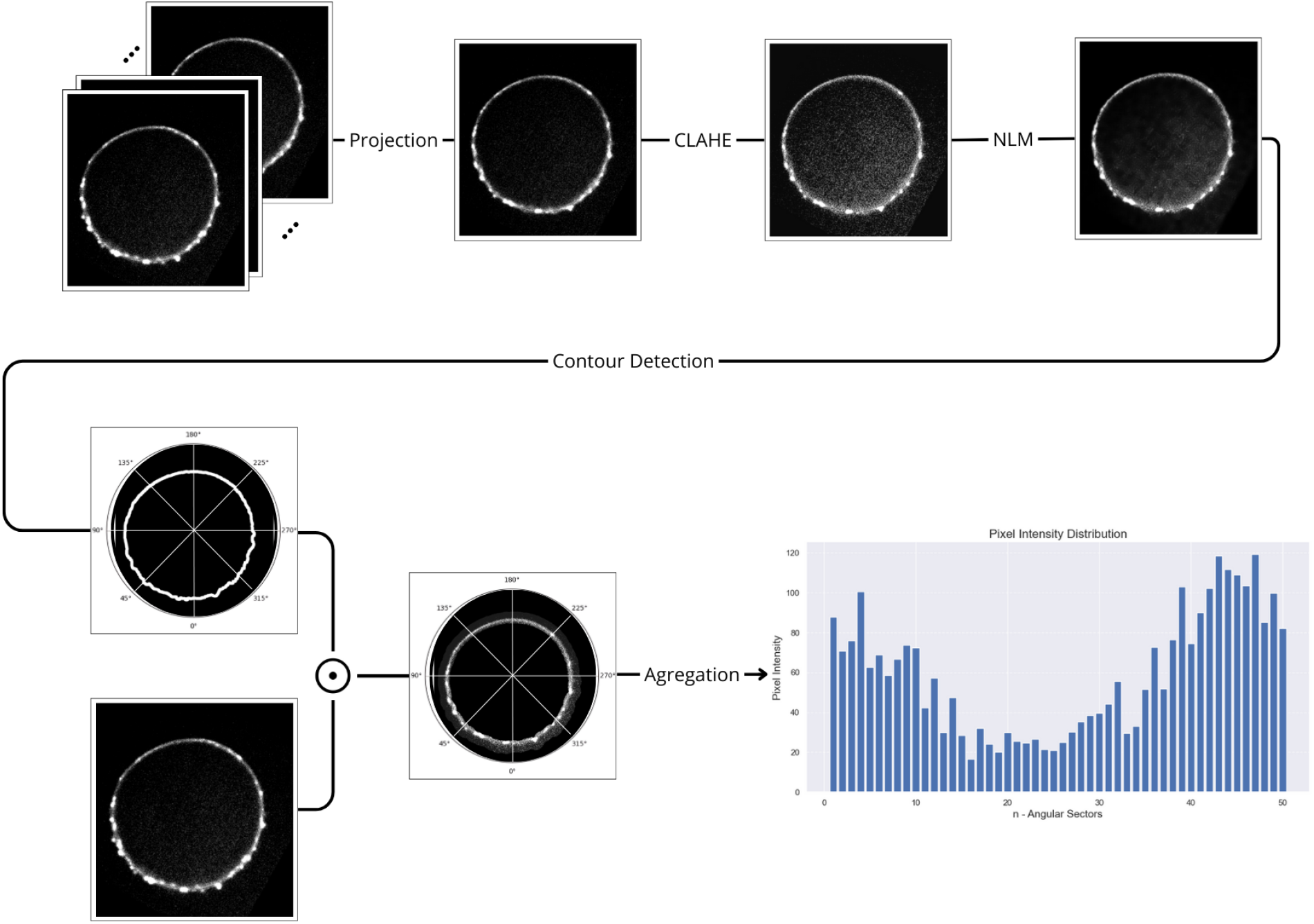
Pipeline of pre-processing videos.

#### 2. Optimal transport

For each frame, we consider an intensity distribution *x_t_* ∈ R*^n^*, where *t* ∈ J1*, n*_frames_). To define the optimal transport formulation for this problem, we introduce two matrices: a coupling matrix *P_t_* ∈ R*^n^*^×^*^n^* and a cost matrix *M* ∈ R*^n^*^×^*^n^*, which are fundamental to the transport optimization framework [5, 6].

(*P_t_*)*_ij_* represents the amount of intensity transported from *i*-th angular sector of the current frame *x_t_*to *j*-th sector of the next frame *x_t_*_+1_. By summing over the columns of *P_t_*, we obtain the intensity distribution of the current frame *x_t_*, while summing over the rows yields the intensity distribution of the next frame *x_t_*_+1_.

The fixed cost matrix *M* (Supplementary Text Fig. 3) defines the transport cost between sectors across consecutive frames. Each element *M_ij_* represents the cost associated with the intensity of the movement from sector *i* ∈ J1*, n*) in the current frame *x_t_* to sector *j* ∈ J1*, n*) in the next frame *x_t_*_+1_, ensuring a physically meaningful transport model [7, 8].

To determine the optimal coupling matrix, we define the transportation polytope *U* (*x_t_, x_t_*_+1_), which constrains the set of feasible coupling matrices:

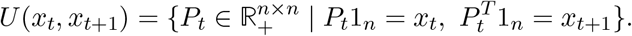

Given this formulation, the optimal transport problem is expressed as finding the coupling matrix *P* ^∗^ that minimizes the total transport cost:

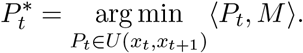

This formulation enables an efficient and principled approach to tracking the evolution of intensity over time. The velocity field is computed by averaging the distance traveled by each element and normalizing by the timestep Δ*t* :

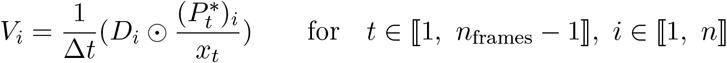

where *D* ∈ R*^n^*^×^*^n^* represents the distance between pixels, incorporating periodicity in movement along the membrane via a clockwise path, as illustrated in Supplementary Text Fig. 3.

**Supplementary Text Figure 3.**
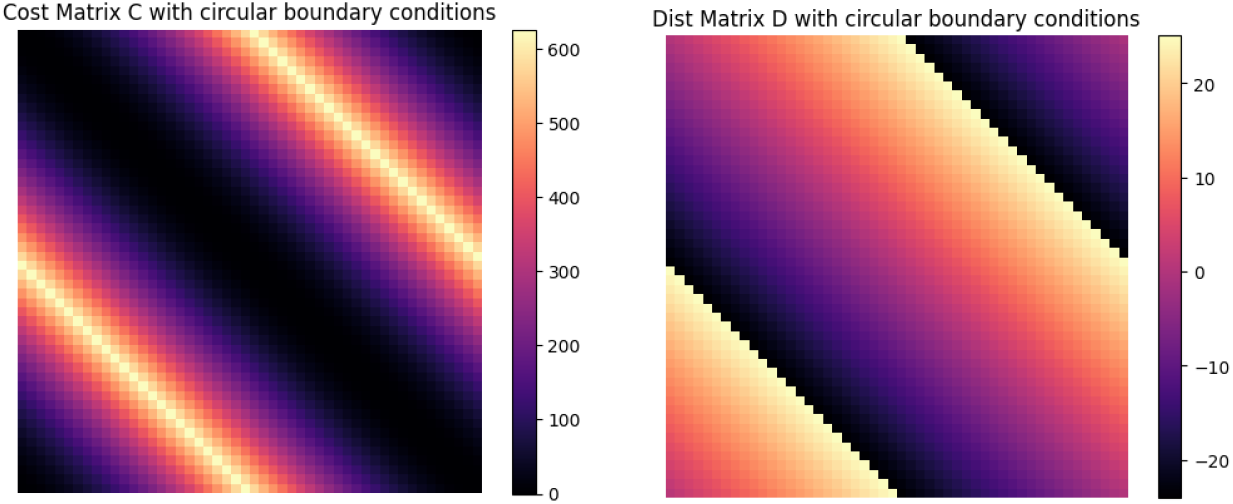
(Left) Cost Matrix, (Right) Distance Matrix.

Solving this optimization problem using linear programming has a complexity of *O*(*n*^3^). Although this is feasible for *n* = 50, scalability becomes a concern for larger numbers of sectors. Additionally, we identified three key challenges with our formulation for determining the optimal coupling matrix:

1. **Computational Cost**: Although our current implementation is tractable, the approach may become impractical for larger-scale problems.
2. **Stability Issues**: The optimization can exhibit numerical instabilities, reducing robustness in certain scenarios.
3. **Intensity Conservation Assumption**: The formulation implicitly assumes conservation of pixel intensities across sectors, which contradicts experimental observations (Supplementary Text Fig. 4). Specifically, the sum of pixel intensities across all sectors is not preserved between consecutive frames.

**Supplementary Text Figure 4.**
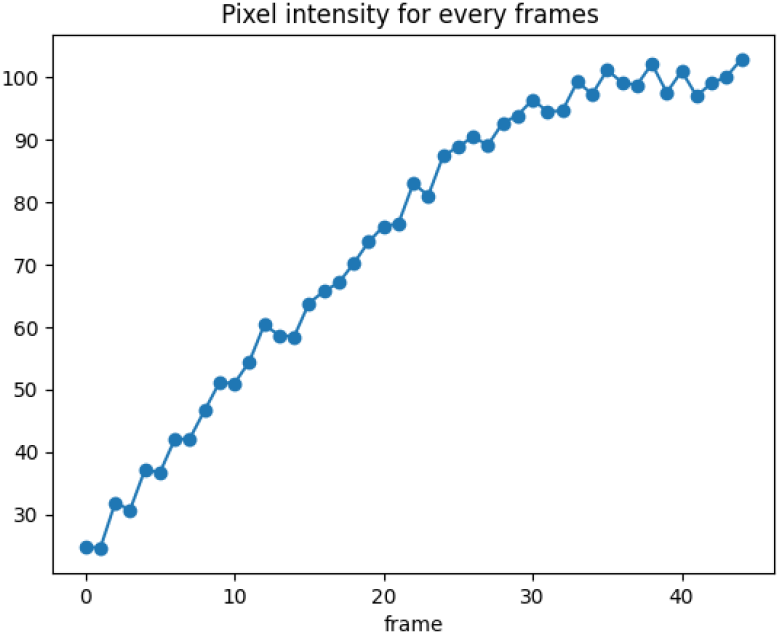
Intensity distribution.

To address these challenges, we take another formulation in which a regularization is introduced into the formulation of the optimal coupling problem, ensuring improved numerical stability and better adherence to empirical intensity distributions.

To address our problem, we introduce the following regularized formulation, incorporating an additional entropy term:

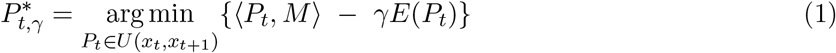

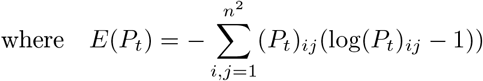

In the following, we introduce an iterative algorithm to solve this regularized problem using Sinkhorn’s algorithm [6]. We begin with the following proposition.

**Proposition 1.** : If *P* ^∗^ = arg min {⟨*P, M* ⟩ − *γE*(*P* )}, then there exists a unique *u, v* ∈ R*^n^* such that

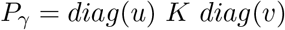

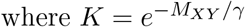

**Proof** [6] : We introduce the Lagrangian multipliers *α, β* ∈ R*^n^* and define the Lagrangian function:

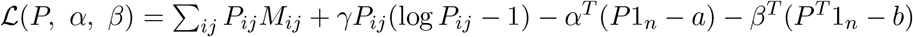

Differentiating with respect to *P_ij_*, we obtain :

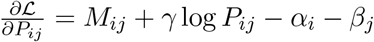

Solving for *P_ij_*, we get

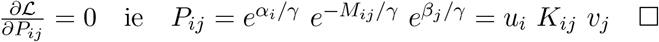

Now, to explicitly determine *u* and *v*, we use the constraints *P_γ_* ∈ *U* (*a, b*), leading to:

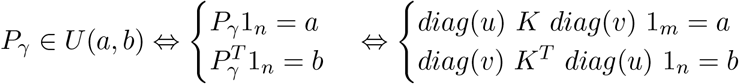

This can be rewritten in component-wise form as :

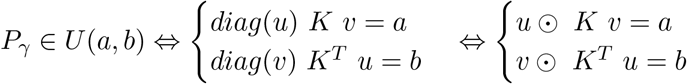

From this, we derive the iterative Sinkhorn [6] after randomly initializing *v*:

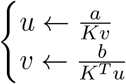

## A hydrodynamic model linking cortical polarization to cytoplasmic reorganization in cRhoA oocytes

In this document, we describe the computational model of cytoplasmic streaming in cRhoA oocytes, its calibration procedure and supplementary results. The computational model specialises our recent framework for modelling coupled surface-bulk viscous flows in animal cells [9] and strikes a good balance between biophysical complexity and accuracy, adopting the necessary simplifications for the sake of its numerical approximation, while still reproducing the experimental behaviour. For the sake of brevity, we will focus on describing the modifications carried out in this work to the original model, in particular, to represent the spindle and its movement. We refer to [9] for full details on the model and its numerical approximation.

### 1. Computational moDel

#### 1.1. Mechanical model of coupled cortex-cytoplasm viscous flows

We consider the system of coupled surface-bulk viscous flow equations given in [9]. The surface phase Γ represents the cell cortex, while the bulk phase Ω represents the cytoplasm. As a first approximation, both are modelled as homogeneous viscous fluids. Flows are driven by a Marangoni-like effect on the cortex, in which gradients of cortical tension are regulated by a diffusing molecular species, representing the myosin-II motors. The cortex is thus modelled as an active fluid surface that drags the enclosing and passive cytoplasmic fluid through a no-slip condition.

The adimensional system of governing equations is given by:

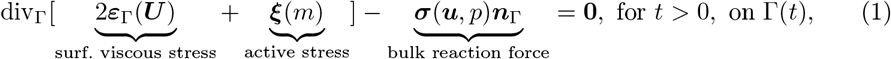

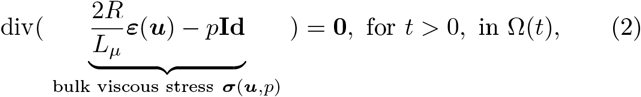

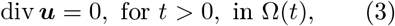

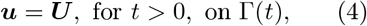

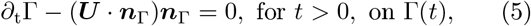

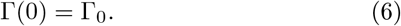

Equations (1-6) are posed in the time-evolving region occupied by the cortex Γ(*t*) and the cytoplasm Ω(*t*); where *t* stands for the adimensional time, *∂_t_* denotes the time derivative and Γ_0_ the initial position of the cortex. ***U*** and *m* represent the surface velocity and myosin-II concentration fields, while ***u*** and *p* the bulk velocity and pressure fields. div_Γ_ and div are the surface and bulk divergence operators, while ***ε***_Γ_ and ***ε*** are the surface and bulk strain rate tensors. *R* is the radius of the cell, *L_µ_* is the surface-to-bulk shear viscosity ratio (a hydrodynamic length scale) and ***n***_Γ_ is the outwards unit normal vector.

Surface stresses in Equation (1) are the resultant of combining the viscous effects 2***ε***_Γ_(***U*** ) with active surface tension ***ξ***(*m*). The latter is governed by the adimensional concentration of myosin-II *m* on the cortex according to the law

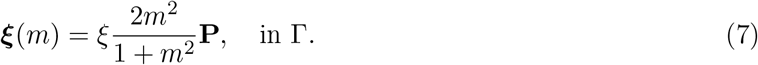

Equation (7) models active tension as a monotonously increasing and saturating Hill function of *m*, scaled by an adimensional coefficient *ξ*, measuring myosin-II activity, where **P** = **Id** − ***n***_Γ_ ⊗ ***n***_Γ_ is the orthogonal projection operator. In contrast to the original system in [9], here the myosin-II concentration *m* is a known field, calibrated experimentally as detailed in Section **1**.**4**.

To conclude, the governing equations are adimensionalised with the following characteristic scales: the initial cell radius *R* and the typical migration time *τ_S_* of the spindle (from the cell centre to the future site of division). According to experimental measurements *R* ≈ 40 µm and spindle migration takes place between BD+4h (BD: nuclear envelope breakdown) and just before anaphase, hence *τ_S_* ≈ 4h. We set *L_µ_/R* ≈ 10^5^ considering typical 3D viscosity and geometrical scales of the cortex and the cytoplasm, according to estimates in [9] from data in previous literature [10, 11].

#### 1.2. Numerical approximation with the finite element method

We discretise the axially symmetric version of Equations (1-6) with 2D *unfitted* finite element methods [12–14]: a family of methods well-suited for problems with moving boundaries and interfaces, as is the case here. In particular, the surface equations are discretised with the Trace FEM [15, 16] and the bulk equations with the Aggregated FEM [17, 18]. Both methods are endowed with mathematical properties that ensure numerical stability and robustness of the discretisation of Equations (1-6).

Numerical implementation was carried out using the Gridap.jl [19–21] finite element software ecosystem, written in the Julia programming language [22]. The code is available at the GitHub repository OocyteAndSpindle (link to Zenodo archive) and supplemented with code demonstrators.

#### 1.3. Representation of the spindle as a frictionless permeated body

The spindle forms a flexible and permeable structure in the cytoplasm, more crowded than other parts of the cytoplasm. A natural way to represent it in the model is as a higher-viscosity region immersed in the cytoplasm. In this sense, the limiting case of a rigid solid-like obstacle has been the choice in other models of cytoplasmic streaming in mouse oocytes [23, 24]. These models examine steady-state flows on a fixed oocyte shape, while our biophysical problem requires accounting for the flow and shape evolution in time. Hence, our computational modelling must be significantly more complex. Besides, the main axes of the spindle of cRhoA oocytes are generally not aligned with the rotation axis. This breaks rotational symmetry (Supplementary Text Fig. 5) and imposes a computationally intensive 3D model that has to be reevaluated for every initial location of the spindle. Above all, we could not clearly identify local perturbations of the flow around the spindle in any of the transmitted light images and STICS analyses. Hence, we could not experimentally infer if, and to what extent, the spindle is behaving as a (permeable) obstacle to the flow.

**Supplementary Text Figure 5.**
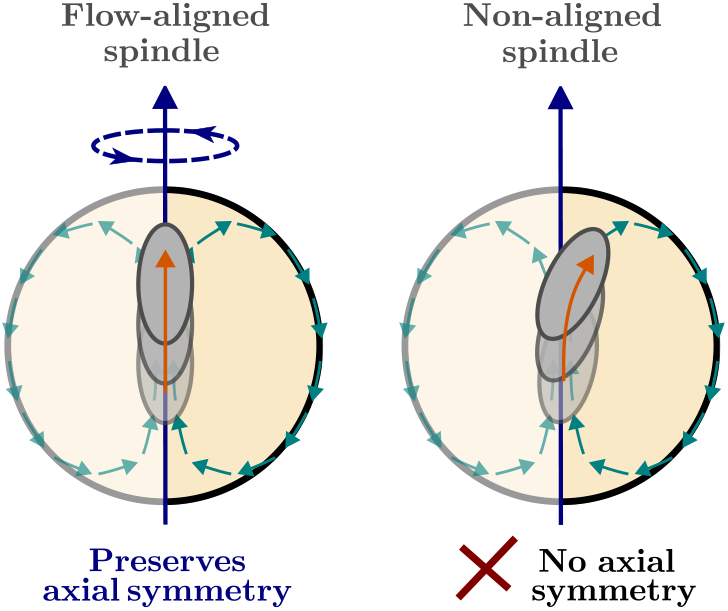
Modelling the spindle as a distinct solid or fluid phase of the cytoplasm breaks rotational symmetry.

Given the computational burden and experimental uncertainty of a solid spindle representation, our modelling assumption considers a different idealised case, where the spindle is still rigid, but not a distinct phase of the cytoplasm. In particular, we assume there is no friction between the spindle and the flow, i.e., the spindle does not perturb the flow at all. It follows that axial symmetry is preserved and the flow is independent of the spindle position. As a result, we can decouple the flow and spindle dynamics in a computationally convenient way, as shown in Supplementary Text Figs. 6A and 6B.

This strategy is divided into two main steps:

1. Calibrate the cytoplasmic flows of the 2D axially symmetric model (see Section 1.4).
2. Compute spindle dynamics as a postprocess of the known 2D velocity field (see Section **1**.**5**).

We will see in the next sections that, despite the simplifying assumptions, our model has enough physical detail to reproduce the experimental observations and describe the hydrodynamics behind the disrupted spindle migration.

#### 1.4. Model calibration

We run numerical simulations from Prophase I (PI) to just before anaphase, i.e., the time interval in which cRhoA polarisation occurs. For this purpose, the model is completed with a calibrated law for the time evolution of the active surface tension from PI to just before anaphase. Recalling Equation 7, the calibration parameters are the time evolution of the myosin-II concentration *m*(*t*) and the activity-scaling coefficient *ξ*.

The experimental calibration is carried out using two sources of quantitative data: (1) time evolution of the oocyte shape from PI to just before anaphase and (2) time-averaged field of cytoplasmic velocities from BD+4h to just before anaphase. The objective is to adjust *m*(*t*) and *ξ* such that the numerical response matches, as much as possible, the cell shape and measured cytoplasmic velocities. At the same time, the calibrated velocities must ensure that a flow-aligned spindle migrates from the oocyte centre to the site of PBE (Polar Body Extrusion) during the last *τ_S_* = 4 physical hours of the simulation.

While the time-averaged velocities are directly provided by the STICS analyses and can be easily compared with the numerical results, shape comparison required further processing of the contour coordinates of the transmitted light images. In particular, we transformed the coordinate data to generate (1) kymographs of radius deviation from the mean oocyte radius (see Fig. 3B) and (2) line plots of oocyte diameter evolution in time (see Supplementary Text Fig. 10). The procedure followed the steps in Supplementary Text Fig. 7.

**Supplementary Text Figure 6.**
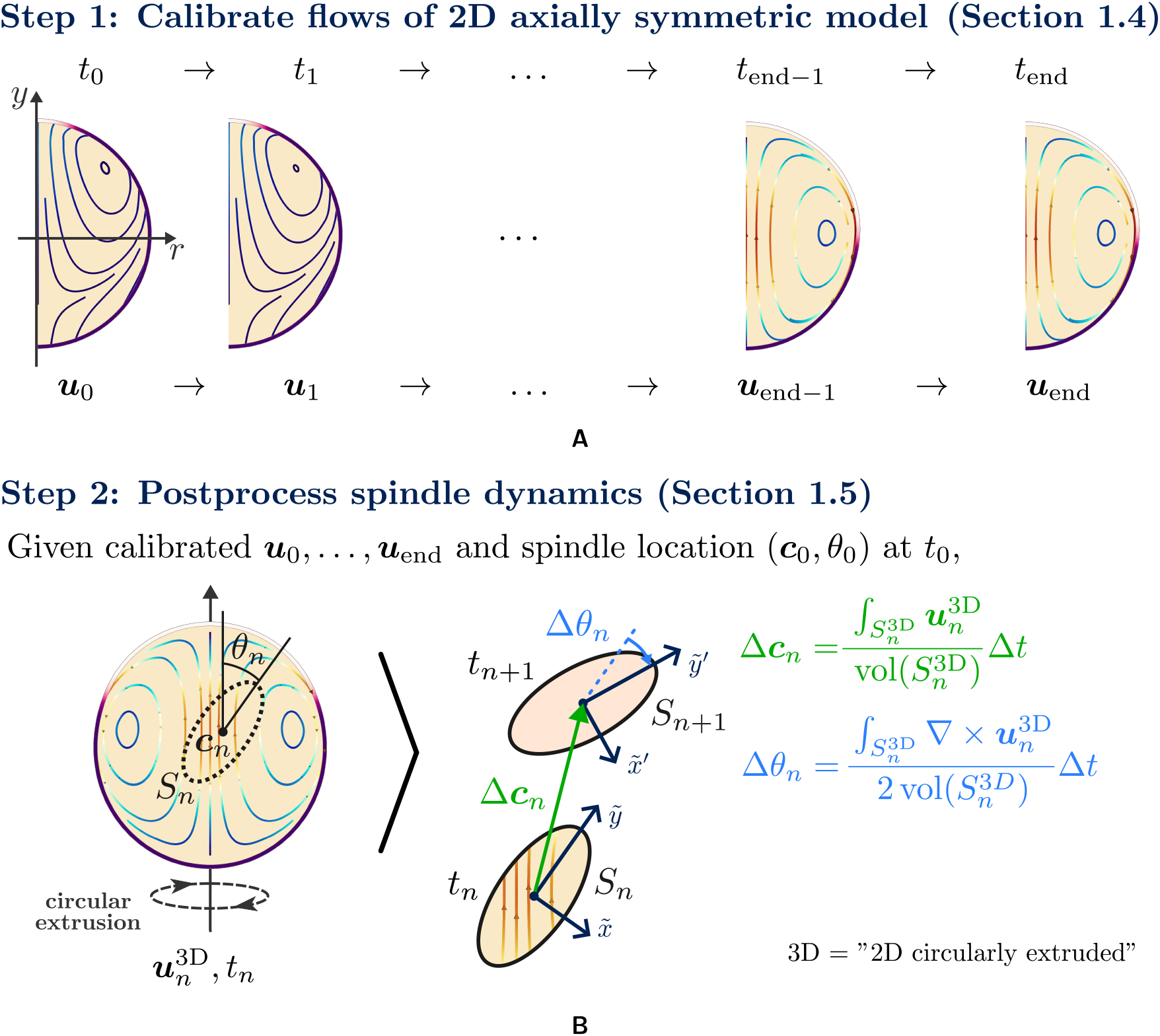
Modelling the spindle as a frictionless permeated body enables decoupling the flow from the spindle dynamics. **A** First, we calibrate the cytoplasmic flows of the 2D axially symmetric model, as described in Section **1**.**4**. **B** Second, the spindle movement at any initial configuration is computed using a circular extrusion of the known 2D velocity field at the cytoplasm, as described in Section **1**.**5**.

Furthermore, a simultaneous adjustment of shape evolution and time-averaged velocities proved to be rather cumbersome. On the other hand, the shape evolution covers the full time window of the physical process and its time evolving nature informs more about the experimental behaviour than the STICS analyses. For this reason, we opted for calibrating the model with the shape evolution data and validating it with the STICS analyses.

### Ansatz for myosin-II evolution over time m(t)

The average kymograph of radius deviation in Fig. 3B reveals the typical pattern of a cortical contraction band, travelling across the oocyte from the future PBE zone to the opposite one. The travelling band emerges spontaneously, probably the result of a mechanochemical instability.

The current model is not able to describe the spontaneous process of symmetry breaking; instead, we calibrate a law for myosin-II evolution over time that produces a similar contraction pattern to the one observed experimentally. This simplification is justified by the fact that the main objective of the model is to describe the hydrodynamics of the cytoplasm and spindle migration, which are driven by the travelling contraction band, rather than its emergence.

If we let the myosin-II concentration in our model evolve as a travelling surface wave, with the back of the crest moving away from the future PBE zone (see Supplementary Text Fig. 8), the response of the active surface stresses precisely corresponds to a surface contraction wave. A similar mechanism is experimentally observed in [25], where RhoA activity coincides with the travelling band of contractility and a wave depleting cortical myosin-II travels from the animal to the vegetal pole.

**Supplementary Text Figure 7.**
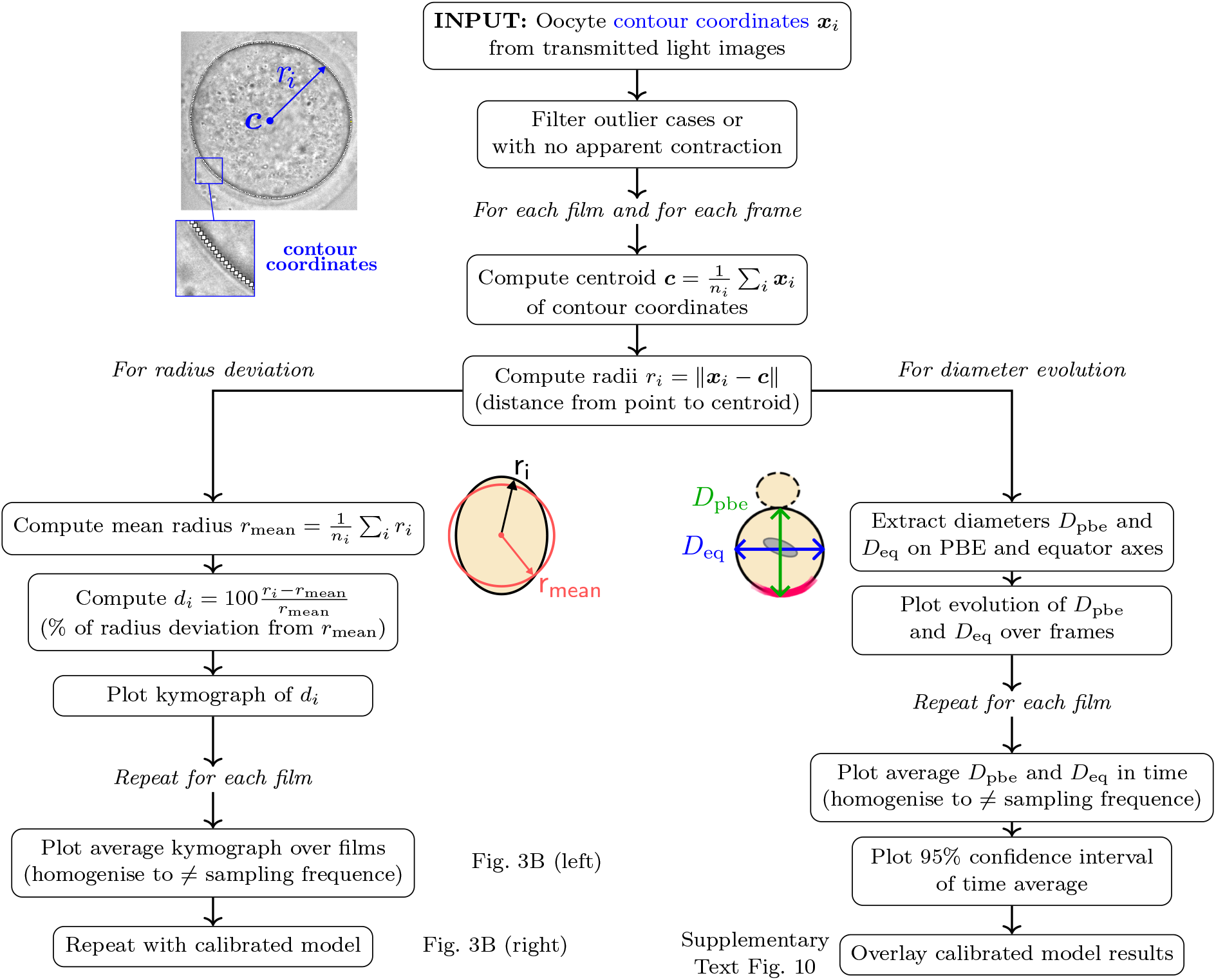
Postprocess to compute radius deviation (left) and diameter evolution (right).

In order to mathematically represent the myosin-II wave, we let

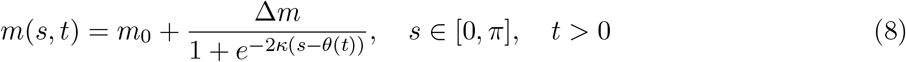

be the ansatz for the evolution of myosin-II over time, where *s* is the arc-length parameter with origin *s* = 0 at the future site of division (*s* = *π* at the opposite side). Equation (8) corresponds to a logistic function, whose minimum is *m*_0_ and maximum is *m*_0_ + Δ*m*; *m*_0_ represents a basal myosin-II density (at the trough of the wave), while Δ*m* measures the jump in myosin-II density at the wave front. *κ* is the steepness of the logistic curve and *θ*(*t*) is the arc-length position of the wave front in time, with *∂_t_ θ*(*t*) representing the speed of the moving front *θ*.

**Supplementary Text Figure 8.**
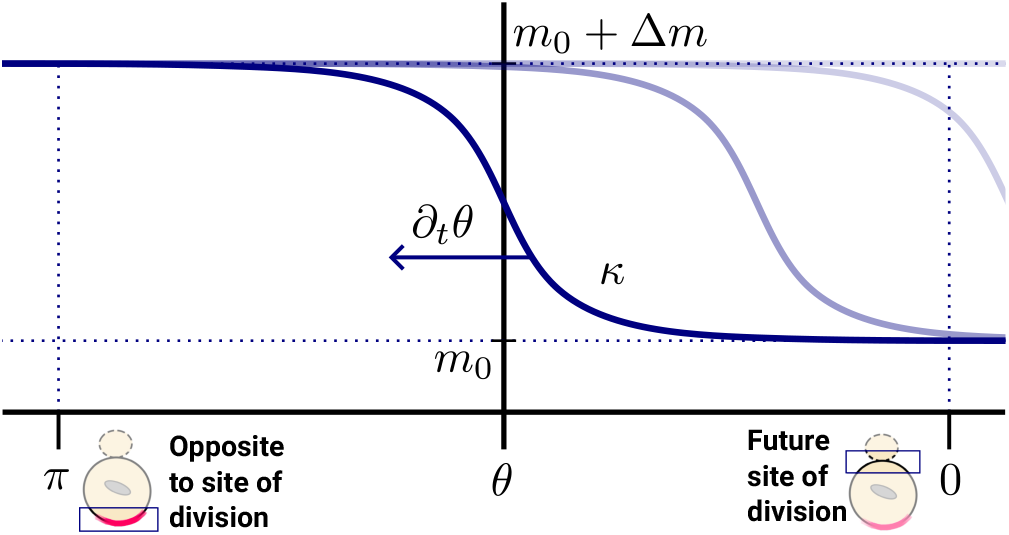
The logistic function in Equation (8) represents myosin-II concentration m over time as a wave travelling across the oocyte surface.

Following manual calibration of Equation (8), our calibrated parameters are *ξ* = 5.0, *m*_0_ = 0.8, Δ*m* = 0.6, *κ* = 10.0 and *θ*(*t*) = 0.52*π/*(1 + *e*^−8(^*^t^*^−0^.^5)^) to meet our objective. We found the most sensitive parameter was the position of the back of the front in time, that is *θ*(*t*), for which the diameter analysis of Supplementary Text Fig. 10A suggests a nonlinear evolution that we tuned with another logistic function.

#### 1.5. Postprocess of spindle dynamics

After calibrating against experiments a *single* 2D axially symmetric FE model, the dynamics of the spindle can be simply computed as a postprocess of the time sequence of the discrete flow velocities {***u***_0_, ***u***_1_*, … ,* ***u***_end−1_, ***u***_end_}.

The procedure is as follows (see Supplementary Text Fig. 6B). Letting ***c****_n_* denote the position of the centre of the spindle and θ*_n_* denote the angle with respect to the axis of rotation at *t* = *t_n_*,

1. compute the average displacement Δ***c****_n_* and rotation angle Δθ*_n_* of the flow particles inside the region occupied by the spindle during the *n*-th time increment,

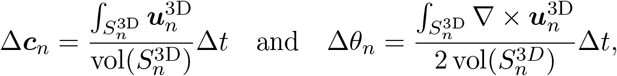

where *S_n_*^3*D*^ and ***u****_n_*^3D^ refer to the 3D spindle surface and the circularly-extruded cytoplasmic velocities at *t* = *t_n_*,

1. update the spindle centre and angle as ***c****_n_*_+1_ ← ***c****_n_* + Δ***c****_n_* and θ*_n_*_+1_ ← θ*_n_* + Δθ*_n_*, that is, apply the rigid motion composed of a translation of magnitude Δ***c****_n_* and a rotation of magnitude Δθ*_n_*,
2. repeat 1. and 2. for the next time increment.

#### 1.6. Assessing spindle rotation across a broad range of initial conditions

We carried out an exhaustive parametric exploration of the model, varying the two key parameters modulating spindle rotation (Supplementary Text Fig. 9): (1) the angle *θ* between the direction of spindle migration and the long axis of the spindle when it is bipolar at BD+4h and (2) the horizontal distance ***d*** to the PBE axis of the spindle centre when it is bipolar. Note that the axis of axial symmetry coincides with the PBE axis and the main direction of the flow in the cytoplasm. Hereafter, we refer to (1) as the *initial spindle angle* and to (2) as the *horizontal offset*.

**Supplementary Text Figure 9.**
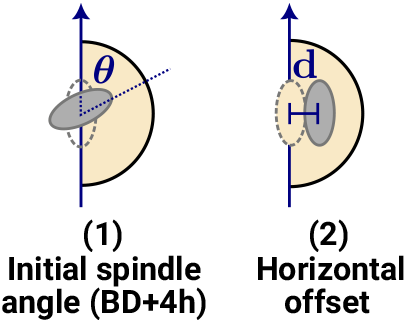
Key initial parameters regulating spindle rotation.

We evaluated the 2D axisymmetric model over all possible initial configurations of the spindle in the (positive) semiplane of axial symmetry (Fig. 3D). Accordingly, the initial spindle angle varies between −*π* and *π*, while the horizontal offset is always positive and varies between 0% and 15% of the oocyte radius (the vast majority of experimental cases fall in this range). Any other 3D spindle configuration can be mapped to the former ones by applying axial symmetry: for instance, spindle configurations with a negative horizontal offset are mirror symmetric to the positive ones.

The bipolar spindle was represented as a smooth spheroid-like shape with a long axis of length 30 µm, a short axis of 14 µm and flattened poles. The implicit function that represents this shape in µm is given by (*y/*15)^2^ +(*x*^2^ +*xz* +*z*^2^)*/*7^2^ = 1. For simplicity, the model assumes the spindle is already bipolar at the beginning, that is, at PI; this simplification has little to no effect in the results, because the numerical spindle barely moves until BD+4h, as imposed by the calibrated Equation (8). Finally, for comparison with experimental results, we computed the initial and variation of spindle migration angles of cRhoA and control oocytes using Fiji’s geometrical analysis tools.

## 2. Supplementary results

### 2.1. Cortical flows driving cRhoA polarity generate a travelling contraction band

The postprocess of the oocyte contour coordinates (Supplementary Text Fig. 7) quantifies the surface contraction wave detected in the transmitted light images. Supplementary Text Fig. 10 shows the time evolution of the deviation from mean oocyte diameter of the diameters at the cross-sections of the PBE and equator axes. The experimental measurements (Supplementary Text Fig. 10A) reconfirm the presence of the cortical contraction wave, as described in Fig. 3B; cRhoA oocytes consistently develop a contraction (negative deviation) at the equator that is mirrored by an expansion at the PBE axis (positive deviation). Maximum contraction/expansion is around 3% as in the kymograph of radius deviation (see Fig. 3B). Continuous lines represent the average time evolution at the PBE and equator axes; they were chosen as the baseline for calibrating the model (Equation (8)). Our numerical model (dashed lines) qualitatively reproduces the temporal pattern observed experimentally, despite difficulties to calibrate when approaching the time of PBE, as PBE onset may be interfering with the oocyte shape (and this is not accounted for in the model). We also report the same quantities for control oocytes in Supplementary Text Fig. 10B to check that, in this case, the diameters of the PBE and equator axes barely vary from PI to just before anaphase. This adds to the evidence that the surface contraction wave during meiosis I is a distinctive trait of cRhoA oocytes.

**Supplementary Text Figure 10.**
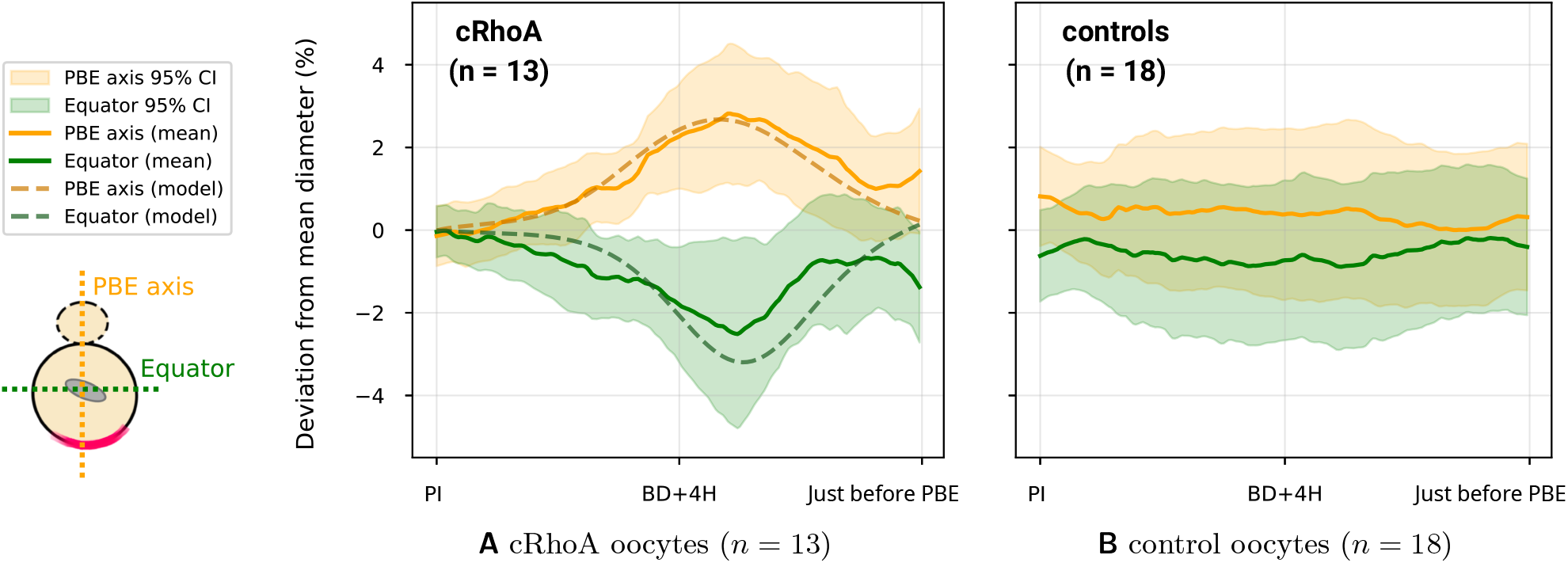
Line plots of diameter evolution in time expose that the cortical contraction wave is a characteristic phenotype of cRhoA oocytes. **A** cRhoA oocytes consistently exhibit a contraction (negative deviation) at the equator, mirrored by an expansion at the PBE axis (positive deviation). **B** Meanwhile the diameters of control oocytes remain stable. Our model (dashed lines) qualitatively reproduces the typical diameter evolution of cRhoA oocytes (continuous lines).

### 2.2. The numerical model captures the observed time-averaged cytoplasmic velocities

Supplementary Text Figs. 11A and 11B report (1) maximum and average velocity magnitudes and (2) average velocity values per component for STICS analyses of controls and cRhoA oocytes. We rotated all cRhoA movies to align the direction of the flow at the oocyte centre with the vertical axis. cRhoA oocytes exhibit a clear distinct phenotype compared to controls, with higher average vertical velocities, as well as higher average and maximum velocity magnitudes, which are consistent with the presence of cytoplasmic streaming.

**Supplementary Text Figure 11.**
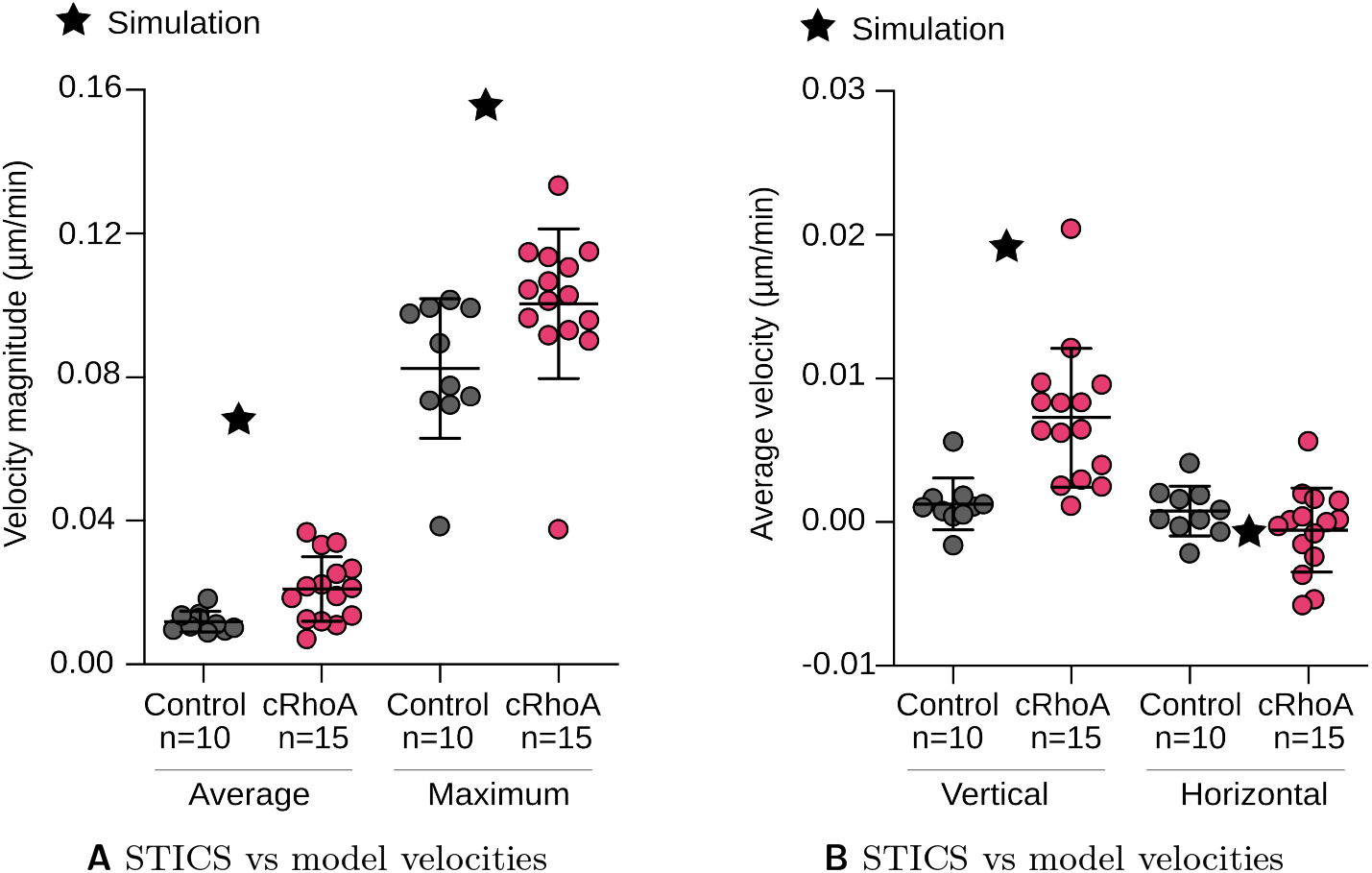
The time-averaged velocity field in the cytoplasm of the calibrated model shows good qualitative agreement with experiments, although it overestimates the velocity magnitudes. This is a consequence of preferring to adjust the shape evolution over the time-averaged velocities (see discussion at the beginning of Section **1**.**4**).

Nonetheless, we observe that the model overestimates the time-averaged cytoplasmic velocities. This follows from the fact that we were not able to calibrate the shape evolution and velocity data at the same time, choosing to calibrate against the shape evolution and use the STICS analyses for validation. Besides, for some STICS analyses, cytoplasmic streaming was not captured in the whole oocyte plane (e.g., only in one half of the oocyte). This may contribute to underestimating experimental average velocities and cause, in part, the discrepancy between the experiments and the model.

### 2.3. The sense of spindle rotation is correctly predicted by the model

Our model predicts that the spindle rotates clockwise at almost any point of the parametric space (see Fig. 3D). An intuitive argument justifying this outcome can be deduced from the spatial 3D pattern of cytoplasmic flows (see Supplementary Text Fig. 12). First, we recall that the parametric space only considers positive horizontal offsets in the semiplane of axial symmetry. Next, we observe that cytoplasmic velocities gradually decrease away from the axis of symmetry. Now, if we consider representative spindle orientations and overlay the flow velocities on selected points, we see that points closer to the axis of symmetry tend to generate the largest torques on the spindle and these torques are, in most cases, clockwise. The same argument implies that the variation of migration angle should increase with the horizontal offset, as predicted by our model.

**Supplementary Text Figure 12.**
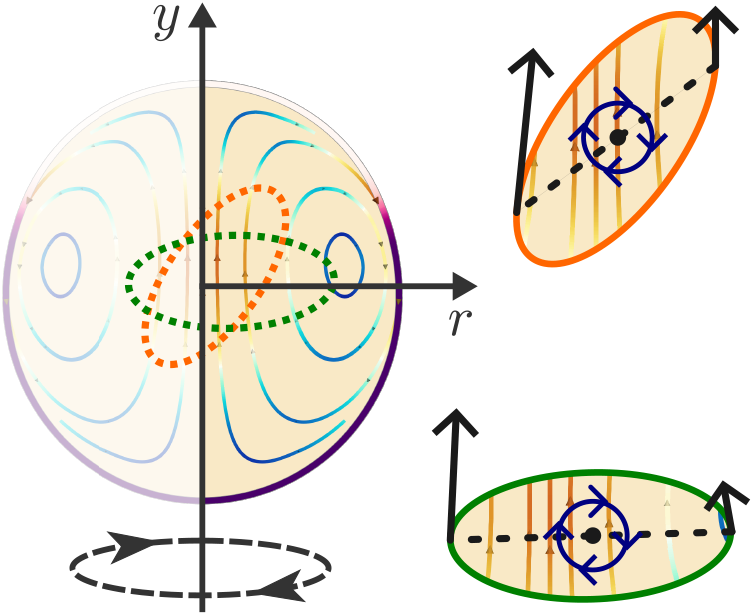
Deducing spindle rotation sense for two cases with positive horizontal offset. Mirror symmetry holds for negative offsets.

## 3. Discussion

Cytoplasmic streaming in cRhoA oocytes is driven by cortical flows that, emerging from a cRhoAinduced instability, establish cortical cRhoA polarity. This study numerically examines the phenomenon under the lens of a minimal mechanical model of coupled cortex-cytoplasm viscous flows.

Numerical model calibration required postprocessing the oocyte shape evolution; the quantitative results add to the body of evidence for the presence of a wave of cortical contractility accompanying the flows. In particular, we show the spatiotemporal patterns of oocyte radii and principal axes lengths (PBE and equator) resemble those identified in, e.g., surface contraction waves of starfish oocytes [25, 26]. Our calibrated model successfully reproduces the flow pattern and shape dynamics of the oocyte and provides a compelling mechanistic explanation for the process. There are two key mechanisms in action: (1) gradients of cortical tension regulated by myosin-II, governing the cortical flows and deformation, and (2) cortical flows dragging cytoplasmic fluid from beneath that recirculates back through the oocyte centre.

In addition, modelling the spindle as a frictionless permeated body drifting in the cytoplasm illustrates the destabilising effect of cytoplasmic streaming on spindle migration. A parametric study of the model lays bare that cytoplasmic flows exert rotational forces on slightly eccentric and/or misoriented spindles; the forces amplify these small perturbations and prevent the spindle from reaching the future site of division with the expected orientation. Qualitative agreement with experiments hints that canonical spindle migration is highly impaired by cytoplasmic streaming.

Our computational model reads as a first approximation of cytoplasmic streaming mediated by cortical contraction waves. It has the minimum ingredients to demonstrate the essence of the active hydrodynamics at play in a simple, yet conclusive way. Further theoretical and computational modelling could elucidate two key aspects of the process. First, all experimental and numerical evidence indicates that the surface contraction wave is spontaneously generated as a mechanochemical instability, such as the polar instability observed in cancer cell migration [27]. The current model cannot capture such type of instabilities, probably due to missing ingredients in the cortex equations. Second, increasing the fidelity in representing the cytoplasm material and the spindle, e.g., as poroviscous materials [28], may quantitatively improve our predictions for cytoplasmic velocities and spindle dynamics, as well as delimit the validity of the common modelling assumption of a viscous cytoplasm.

## Notes

### Competing Interest Statement

The authors have declared no competing interest.

## References

1. Nikalayevich, E., Zollo, N. & Verlhac, M.-H. Impact of organelle architecture on oocyte developmental potential. Current Opinion in Cell Biology 95, 102556 (2025).

2. Jentoft, I. M. A. & Schuh, M. Protein Storage in Oocytes: Implications for Oocyte Quality, Embryonic Development, and Female Fertility. Annu Rev Cell Dev Biol 41, 15–43 (2025).

3. Harris, P. J., Clason, E. L. & Prier, K. R. Tubulin polymerization in unfertilized sea-urchin eggs induced by elevated temperature. J Cell Sci 93, 9–17 (1989).

4. Gard, D. L. Confocal microscopy and 3-D reconstruction of the cytoskeleton of Xenopus oocytes. Microscopy Research and Technique 44, 388–414 (1999).

5. Miyazaki, A., Kamitsubo, E. & Nemoto, S.-I. Premeiotic Aster as a Device to Anchor the Germinal Vesicle to the Cell Surface of the Presumptive Animal Pole in Starfish Oocytes. Developmental Biology 218, 161–171 (2000).

6. Miyazaki, A., Kato, K. H. & Nemoto, S. Role of microtubules and centrosomes in the eccentric relocation of the germinal vesicle upon meiosis reinitiation in sea-cucumber oocytes. Developmental Biology 280, 237–247 (2005).

7. Amiel, A. & Houliston, E. Three distinct RNA localization mechanisms contribute to oocyte polarity establishment in the cnidarian *Clytia hemisphærica*. Developmental Biology 327, 191–203 (2009).

8. McNally, K. L., Martin, J. L., Ellefson, M. & McNally, F. J. Kinesin-dependent transport results in polarized migration of the nucleus in oocytes and inward movement of yolk granules in meiotic embryos. Developmental biology 339, 126 (2009).

9. Elkouby, Y. M., Jamieson-Lucy, A. & Mullins, M. C. Oocyte Polarization Is Coupled to the Chromosomal Bouquet, a Conserved Polarized Nuclear Configuration in Meiosis. PLOS Biology 14, e1002335 (2016).

10. Tissot, N. et al. Distinct molecular cues ensure a robust microtubule-dependent nuclear positioning in the Drosophila oocyte. Nature Communications 8, 15168 (2017).

11. Tokuhisa, M., Muto, M. & Nishida, H. Eccentric position of the germinal vesicle and cortical flow during oocyte maturation specify the animal-vegetal axis of ascidian embryos. Development 144, 897–904 (2017).

12. Almonacid, M., Terret, M.-E. & Verlhac, M.-H. Control of nucleus positioning in mouse oocytes. Semin Cell Dev Biol 82, 34–40 (2018).

13. Halet, G. & Carroll, J. Rac Activity Is Polarized and Regulates Meiotic Spindle Stability and Anchoring in Mammalian Oocytes. Developmental Cell 12, 309–317 (2007).

14. FitzHarris, G., Marangos, P. & Carroll, J. Changes in endoplasmic reticulum structure during mouse oocyte maturation are controlled by the cytoskeleton and cytoplasmic dynein. Developmental Biology 305, 133–144 (2007).

15. Dalton, C. M. & Carroll, J. Biased inheritance of mitochondria during asymmetric cell division in the mouse oocyte. Journal of Cell Science 126, 2955–2964 (2013).

16. Almonacid, M. et al. Active diffusion positions the nucleus in mouse oocytes. Nat Cell Biol 17, 470–479 (2015).

17. Verlhac, M.-H., Lefebvre, C., Guillaud, P., Rassinier, P. & Maro, B. Asymmetric division in mouse oocytes: with or without Mos. Current Biology 10, 1303–1306 (2000).

18. Azoury, J. et al. Spindle Positioning in Mouse Oocytes Relies on a Dynamic Meshwork of Actin Filaments. Current Biology 18, 1514–1519 (2008).

19. Schuh, M. & Ellenberg, J. A New Model for Asymmetric Spindle Positioning in Mouse Oocytes. Current Biology 18, 1986–1992 (2008).

20. Chaigne, A. et al. A soft cortex is essential for asymmetric spindle positioning in mouse oocytes. Nat Cell Biol 15, 958–966 (2013).

21. Bennabi, I. et al. Artificially decreasing cortical tension generates aneuploidy in mouse oocytes. Nat Commun 11, 1649 (2020).

22. Chaigne, A. et al. A narrow window of cortical tension guides asymmetric spindle positioning in the mouse oocyte. Nat Commun 6, 6027 (2015).

23. Nikalayevich, E. et al. Aberrant cortex contractions impact mammalian oocyte quality. Developmental Cell 59, 841–852.e7 (2024).

24. Bulteau, R. et al. Atomic Force Microscopy Reveals Differences In Mechanical Properties Linked To Cortical Structure In Mouse And Human Oocytes. Small 21, e2500221 (2025).

25. Longo, F. J. & Chen, D.-Y. Development of cortical polarity in mouse eggs: Involvement of the meiotic apparatus. Developmental Biology 107, 382–394 (1985).

26. Maro, B., Johnson, M. H., Webb, M. & Flach, G. Mechanism of polar body formation in the mouse oocyte: an interaction between the chromosomes, the cytoskeleton and the plasma membrane. J Embryol Exp Morphol 92, 11–32 (1986).

27. Brunet, S. & Verlhac, M. H. Positioning to get out of meiosis: the asymmetry of division. Human Reproduction Update 17, 68–75 (2011).

28. Miyado, K. et al. Requirement of CD9 on the Egg Plasma Membrane for Fertilization. Science 287, 321–324 (2000).

29. Bianchi, E., Doe, B., Goulding, D. & Wright, G. J. Juno is the egg Izumo receptor and is essential for mammalian fertilisation. Nature 508, 483–487 (2014).

30. Mori, M. et al. RanGTP and the actin cytoskeleton keep paternal and maternal chromosomes apart during fertilization. Journal of Cell Biology 220, e202012001 (2021).

31. Zhang, X. et al. Polar Body Emission Requires a RhoA Contractile Ring and Cdc42- Mediated Membrane Protrusion. Developmental Cell 15, 386–400 (2008).

32. Dehapiot, B., Carrière, V., Carroll, J. & Halet, G. Polarized Cdc42 activation promotes polar body protrusion and asymmetric division in mouse oocytes. Dev Biol 377, 202–212 (2013).

33. Wang, Z.-B. et al. Specific deletion of Cdc42 does not affect meiotic spindle organization/migration and homologous chromosome segregation but disrupts polarity establishment and cytokinesis in mouse oocytes. MBoC 24, 3832–3841 (2013).

34. Yuen, W. S. et al. Polo-like kinase 1 promotes Cdc42-induced actin polymerization for asymmetric division in oocytes. Open Biol 13, 220326 (2023).

35. Shihabi, A., Bulteau, R., Barbier, L. & Terret, M.-E. From Cytoskeletal Remodeling to Oocyte Quality: The Emerging Role of Mechanics. Advanced Science 13, e76323 (2026).

36. Bodor, D. L., Pönisch, W., Endres, R. G. & Paluch, E. K. Of Cell Shapes and Motion: The Physical Basis of Animal Cell Migration. Dev Cell 52, 550–562 (2020).

37. De Belly, H., Paluch, E. K. & Chalut, K. J. Interplay between mechanics and signalling in regulating cell fate. Nat Rev Mol Cell Biol 23, 465–480 (2022).

38. Gupta, V. & Chaudhuri, O. Mechanical regulation of cell cycle progression and division. Trends Cell Biol 32, 773–785 (2022).

39. Fiore, V. F., Almagro, J. & Fuchs, E. Shaping epithelial tissues by stem cell mechanics in development and cancer. Nat Rev Mol Cell Biol 26, 442–455 (2025).

40. Su, Z., Chen, Z., Ma, K., Chen, H. & Ho, J. W. K. Molecular determinants of intrinsic cellular stiffness in health and disease. Biophys Rev 14, 1197–1209 (2022).

41. Brauer, E. et al. Dissecting the influence of cellular senescence on cell mechanics and extracellular matrix formation in vitro. Aging Cell 22, e13744 (2023).

42. Belousova, P., Kwon, S., Jung, M.-H. & Kim, K. S. Impact of mechanical property alterations on cancer cell motility and metastasis. Critical Reviews in Oncology/Hematology 220, 105165 (2026).

43. Cavanaugh, K. E. et al. Elevated contractility drives age-associated implantation failure in mouse embryos from aged females. Nat Cell Biol 28, 1799–1813 (2026).

44. Yanez, L. Z., Han, J., Behr, B. B., Pera, R. A. R. & Camarillo, D. B. Human oocyte developmental potential is predicted by mechanical properties within hours after fertilization. Nat Commun 7, 10809 (2016).

45. van der Kammen, R. et al. Knockout of the Arp2/3 complex in epidermis causes a psoriasis- like disease hallmarked by hyperactivation of transcription factor Nrf2. Development 144, 4588–4603 (2017).

46. Wang, H. et al. Symmetry breaking in hydrodynamic forces drives meiotic spindle rotation in mammalian oocytes. Sci. Adv. 6, eaaz5004 (2020).

47. Louvet, S., Aghion, J., Santa-Maria, A., Mangeat, P. & Maro, B. Ezrin Becomes Restricted to Outer Cells Following Asymmetrical Division in the Preimplantation Mouse Embryo. Developmental Biology 177, 568–579 (1996).

48. Riento, K. & Ridley, A. J. ROCKs: multifunctional kinases in cell behaviour. Nat Rev Mol Cell Biol 4, 446–456 (2003).

49. De Belly, H. et al. Cell protrusions and contractions generate long-range membrane tension propagation. Cell 186, 3049–3061.e15 (2023).

50. Hawkins, R. J. et al. Spontaneous Contractility-Mediated Cortical Flow Generates Cell Migration in Three-Dimensional Environments. Biophysical Journal 101, 1041–1045 (2011).

51. Ruprecht, V. et al. Cortical Contractility Triggers a Stochastic Switch to Fast Amoeboid Cell Motility. Cell 160, 673–685 (2015).

52. Yi, K. et al. Dynamic maintenance of asymmetric meiotic spindle position through Arp2/3- complex-driven cytoplasmic streaming in mouse oocytes. Nat Cell Biol 13, 1252–1258 (2011).

53. Li, S. et al. Basal actomyosin pulses expand epithelium coordinating cell flattening and tissue elongation. Nat Commun 15, 3000 (2024).

54. Leerberg, D. M., Avillion, G. B., Priya, R., Stainier, D. Y. R. & Yelon, D. Regionalized regulation of actomyosin organization influences cardiomyocyte cell shape changes during chamber curvature formation. Nat Commun 17, 3768 (2026).

55. Colin, A. et al. Active diffusion in oocytes nonspecifically centers large objects during prophase I and meiosis I. Journal of Cell Biology 219, e201908195 (2020).

56. Neiva, E. & Turlier, H. Unfitted finite element modelling of surface-bulk viscous flows in animal cells. Journal of Computational Physics 560, 114926 (2026).

57. Mietke, A., Jemseena, V., Kumar, K. V., Sbalzarini, I. F. & Jülicher, F. Minimal Model of Cellular Symmetry Breaking. Phys Rev Lett 123, 188101 (2019).

58. 58. Johnson, C. Numerical Solution of Partial Differential Equations by the Finite Element Method. (Cambridge [England] ; New York : Cambridge University Press, 1987).

59. Mori, M. et al. RanGTP and the actin cytoskeleton keep paternal and maternal chromosomes apart during fertilization. Journal of Cell Biology 220, e202012001 (2021).

60. Cheeseman, L. P., Boulanger, J., Bond, L. M. & Schuh, M. Two pathways regulate cortical granule translocation to prevent polyspermy in mouse oocytes. Nat Commun 7, 13726 (2016).

61. Letort, G. et al. An interpretable and versatile machine learning approach for oocyte phenotyping. Journal of Cell Science 135, jcs260281 (2022).

62. Crozet, F. et al. Filopodia-like protrusions of adjacent somatic cells shape the developmental potential of oocytes. Life Sci Alliance 6, e202301963 (2023).

63. Carvalho, K. et al. Actin polymerization or myosin contraction: two ways to build up cortical tension for symmetry breaking. Philos Trans R Soc Lond B Biol Sci 368, 20130005 (2013).

64. Munro, E., Nance, J. & Priess, J. R. Cortical Flows Powered by Asymmetrical Contraction Transport PAR Proteins to Establish and Maintain Anterior-Posterior Polarity in the Early *C. elegans* Embryo. Developmental Cell 7, 413–424 (2004).

65. Mayer, M., Depken, M., Bois, J. S., Jülicher, F. & Grill, S. W. Anisotropies in cortical tension reveal the physical basis of polarizing cortical flows. Nature 467, 617–621 (2010).

66. Panda, Y. A. & Fischer-Friedrich, E. Localized actin cortex perturbation generates cell-scale membrane tension gradients. 2026.05.06.721853 Preprint at 10.64898/2026.05.06.721853 (2026).

67. Nishikawa, M., Naganathan, S. R., Jülicher, F. & Grill, S. W. Controlling contractile instabilities in the actomyosin cortex. eLife 6, e19595 (2017).

68. De Belly, H. et al. Long-range mutual activation establishes Rho and Rac polarity during cell migration. Nat Cell Biol 28, 1244–1257 (2026).

69. Galbraith, C. G., English, B. P., Boehm, U. & Galbraith, J. A. Compartmentalized cytoplasmic tradewinds direct soluble proteins. Nat Commun 17, 2589 (2026).

70. Niwayama, R. et al. Bayesian Inference of Forces Causing Cytoplasmic Streaming in Caenorhabditis elegans Embryos and Mouse Oocytes. PLOS ONE 11, e0159917 (2016).

71. Dehapiot, B. et al. RhoA- and Cdc42-induced antagonistic forces underlie symmetry breaking and spindle rotation in mouse oocytes. PLoS Biol 19, e3001376 (2021).

72. Lee, I.-W. et al. Chromatin- and actin-mediated mitochondrial streaming leads to patterning of mitochondrial distribution in oocytes. Nat Commun 17, 6569 (2026).

73. Reynier, P. et al. Mitochondrial DNA content affects the fertilizability of human oocytes. Mol Hum Reprod 7, 425–429 (2001).

74. Santos, T. A., El Shourbagy, S. & St. John, J. C. Mitochondrial content reflects oocyte variability and fertilization outcome. Fertility and Sterility 85, 584–591 (2006).

75. El Shourbagy, S. H., Spikings, E. C., Freitas, M. & St John, J. C. Mitochondria directly influence fertilisation outcome in the pig. Reproduction 131, 233–245 (2006).

76. Ge, H. et al. The importance of mitochondrial metabolic activity and mitochondrial DNA replication during oocyte maturation in vitro on oocyte quality and subsequent embryo developmental competence. Molecular Reproduction and Development 79, 392–401 (2012).

77. Murakoshi, Y. et al. Embryo developmental capability and pregnancy outcome are related to the mitochondrial DNA copy number and ooplasmic volume. J Assist Reprod Genet 30, 1367–1375 (2013).

78. Ducibella, T., Kurasawa, S., Duffy, P., Kopf, G. S. & Schultz, R. M. Regulation of the Polyspermy Block in the Mouse Egg: Maturation-Dependent Differences in Cortical Granule Exocytosis and Zona Pellucida Modifications Induced by Inositol 1,4,5-Trisphosphate and an Activator of Protein Kinase C. Biol Reprod **48**, 1251–1257 (1993).

79. Evans, J. P. Preventing polyspermy in mammalian eggs—Contributions of the membrane block and other mechanisms. Molecular Reproduction and Development 87, 341–349 (2020).

80. Dubois, Y. et al. Roles of the zona pellucida in gamete fusion and of the perivitelline space in blocking polyspermy in mice. EMBO Rep 27, 774–792 (2026).

81. Rozema, D. & Maître, J.-L. Forces Shaping the Blastocyst. Cold Spring Harb Perspect Biol 17, a041519 (2025).

82. Reis, A., Chang, H.-Y., Levasseur, M. & Jones, K. T. APCcdh1 activity in mouse oocytes prevents entry into the first meiotic division. Nat Cell Biol 8, 539–540 (2006).

83. Manil-Ségalen, M. et al. Chromosome structural anomalies due to aberrant spindle forces exerted at gene editing sites in meiosis. J Cell Biol 217, 3416–3430 (2018).

84. Inoue, N., Saito, T. & Wada, I. Unveiling a novel function of CD9 in surface compartmentalization of oocytes. Development dev.189985 (2020) doi:10.1242/dev.189985.

85. Łuksza, M., Queguigner, I., Verlhac, M.-H. & Brunet, S. Rebuilding MTOCs upon centriole loss during mouse oogenesis. Developmental Biology 382, 48–56 (2013).

86. Schindelin, J. et al. Fiji: an open-source platform for biological-image analysis. Nat Methods 9, 676–682 (2012).

87. Hebert, B., Costantino, S. & Wiseman, P. Spatiotemporal Image Correlation Spectroscopy (STICS) Theory, Verification, and Application to Protein Velocity Mapping in Living CHO Cells. Biophysical journal 88, 3601–14 (2005).

## References

[1] K. Zuiderveld. “Contrast Limited Adaptive Histogram Equalization”. In: Graphics Gems IV. Ed. by P. S. Heckbert. Academic Press, 1994, pp. 474–485.

[2] A. Buades, B. Coll, and J.-M. Morel. “A Non-Local Algorithm for Image Denoising”. In: IEEE Computer Vision and Pattern Recognition (CVPR) 2 (2005), pp. 60–65.

[3] N. Otsu. “A Threshold Selection Method from Gray-Level Histograms”. In: IEEE Transactions on Systems, Man, and Cybernetics 9.1 (1979), pp. 62–66.

[4] S. Suzuki and K. Abe. “Topological Structural Analysis of Digitized Binary Images by Border Following”. In: Computer Vision, Graphics, and Image Processing 30.1 (1985), pp. 32–46.

[5] C. Villani. Topics in Optimal Transportation. American Mathematical Society, 2003.

[6] M. Cuturi. “Sinkhorn distances: Lightspeed computation of optimal transport”. In: Advances in Neural Information Processing Systems 26 (2013), pp. 2292–2300.

[7] G. Peyré and M. Cuturi. “Computational optimal transport”. In: Foundations and Trends® in Machine Learning 11.5-6 (2019), pp. 355–607.

[8] J. Solomon. Optimal Transport on Discrete Domains. Cambridge University Press, 2018.

[9] E. Neiva and H. Turlier. “Unfitted finite element modelling of surface-bulk viscous flows in animal cells”. In: Journal of Computational Physics 560 (2026), p. 114926. doi: 10.1016/j.jcp.2026.114926.

[10] H. Turlier et al. “Furrow constriction in animal cell cytokinesis”. In: Biophysical Journal 106.1 (2014), pp. 114–123. doi: 10.1016/j.bpj.2013.11.014.

[11] P. A. Valberg and H. A. Feldman. “Magnetic particle motions within living cells. Measurement of cytoplasmic viscosity and motile activity”. In: Biophysical Journal 52.4 (1987), pp. 551–561. doi: 10.1016/S0006-3495(87)83244-7.

[12] C. S. Peskin. “Flow patterns around heart valves: a numerical method”. In: Journal of computational physics 10.2 (1972), pp. 252–271. doi: 10.1016/0021-9991(72)90065-4.

[13] F. de Prenter et al. “Stability and conditioning of immersed finite element methods: analysis and remedies”. In: Archives of Computational Methods in Engineering 30.6 (2023), pp. 3617–3656. doi: 10.1007/s11831-023-09913-0.

[14] E. Burman et al. “Cut finite element methods”. In: Acta Numerica 34 (2025), 1–121. doi: 10.1017/S0962492925000017.

[15] M. A. Olshanskii and A. Reusken. “Trace Finite Element Methods for PDEs on Surfaces”. In: Geometrically Unfitted Finite Element Methods and Applications. Ed. by S. P. A. Bordas et al. Cham: Springer International Publishing, 2017, pp. 211–258.

[16] C. Lehrenfeld, M. A. Olshanskii, and X. Xu. “A stabilized trace finite element method for partial differential equations on evolving surfaces”. In: SIAM Journal on Numerical Analysis 56.3 (2018), pp. 1643–1672. doi: 10.1137/17M1148633.

[17] S. Badia, F. Verdugo, and A. F. Martín. “The aggregated unfitted finite element method for elliptic problems”. In: Computer Methods in Applied Mechanics and Engineering 336 (2018), pp. 533–553. doi: 10.1016/j.cma.2018.03.022.

[18] S. Badia, A. F. Martin, and F. Verdugo. “Mixed Aggregated Finite Element Methods for the Unfitted Discretization of the Stokes Problem”. In: SIAM Journal on Scientific Computing 40.6 (2018), B1541– B1576. doi: 10.1137/18M1185624.

[19] S. Badia and F. Verdugo. “Gridap: An extensible Finite Element toolbox in Julia”. In: Journal of Open Source Software 5.52 (2020), p. 2520. doi: 10.21105/joss.02520. url: 10.21105/joss.02520.

[20] F. Verdugo and S. Badia. “The software design of Gridap: A Finite Element package based on the Julia JIT compiler”. In: Computer Physics Communications 276 (2022), p. 108341. doi: 10.1016/j.cpc.2022.108341. url: 10.1016/j.cpc.2022.108341.

[21] F. Verdugo et al. *gridap/GridapEmbedded.jl: v0.9.5*. Version v0.9.5. Oct. 2024. doi: 10.5281/zenodo.13949319. url: https://github.com/gridap/GridapEmbedded.jl.

[22] J. Bezanson et al. “Julia: A fresh approach to numerical computing”. In: SIAM Review 59.1 (2017), pp. 65–98. doi: 10.1137/141000671. url: https://epubs.siam.org/doi/10.1137/141000671.

[23] K. Yi et al. “Dynamic maintenance of asymmetric meiotic spindle position through Arp2/3-complex-driven cytoplasmic streaming in mouse oocytes”. In: Nature cell biology 13.10 (2011), pp. 1252–1258. doi: 10.1038/ncb2320.

[24] W. Liao and E. Lauga. “Hydrodynamic Mechanism for Stable Spindle Positioning in Meiosis II Oocytes”. In: PRX Life 2 (4 2024), p. 043003. doi: 10.1103/PRXLife.2.043003.

[25] J. Bischof, et al. “A cdk1 gradient guides surface contraction waves in oocytes”. In: Nature communications 8.1 (2017), p. 849.

[26] N. Klughammer et al. “Cytoplasmic flows in starfish oocytes are fully determined by cortical contractions”. In: PLoS computational biology 14.11 (2018), e1006588. doi: 10.1371/journal.pcbi.1006588.

[27] R. J. Hawkins et al. “Spontaneous contractility-mediated cortical flow generates cell migration in three-dimensional environments”. In: Biophysical Journal 101.5 (2011), pp. 1041–1045. doi: 10.1016/j.bpj.2011.07.038.

[28] A. Mogilner and A. Manhart. “Intracellular fluid mechanics: Coupling cytoplasmic flow with active cytoskeletal gel”. In: Annual Review of Fluid Mechanics 50.1 (2018), pp. 347–370. doi: 10.1146/annurev-fluid-010816-060238.

